# Precision fMRI reveals that the language-selective network supports both phrase-structure building and lexical access during language production

**DOI:** 10.1101/2021.09.10.459596

**Authors:** Jennifer Hu, Hannah Small, Hope Kean, Atsushi Takahashi, Leo Zekelman, Daniel Kleinman, Elizabeth Ryan, Alfonso Nieto-Castañón, Victor Ferreira, Evelina Fedorenko

**Author notes:** Equal contributors (Jennifer Hu, Hannah Small). Corresponding Authors Jennifer Hu or Ev Fedorenko /; 43 Vassar Street, Room 46-3037, Cambridge, MA, 02139.

## Abstract

A fronto-temporal brain network has long been implicated in language comprehension. However, this network’s role in language production remains debated. In particular, it remains unclear whether all or only some language regions contribute to production, and which aspects of production these regions support. Across three fMRI experiments that rely on robust individual-subject analyses, we characterize the language network’s response to high-level production demands. We report three novel results. First, sentence production, spoken or typed, elicits a strong response throughout the language network. Second, the language network responds to both phrase-structure building and lexical access demands, although the response to phrase-structure building is stronger and more spatially extensive, present in every language region. Finally, contra some proposals, we find no evidence of brain regions—within or outside the language network—that selectively support phrase-structure building in production relative to comprehension. Instead, all language regions respond more strongly during production than comprehension, suggesting that production incurs a greater cost for the language network. Together, these results align with the idea that language comprehension and production draw on the same knowledge representations, which are stored in a distributed manner within the language-selective network and are used to both interpret and generate linguistic utterances.

## Main text

Although the scientific enterprise of language neuroscience began with a report about an aspect of language *production* (articulatory abilities; Broca, 1861), the field has been largely dominated by investigations of language *comprehension*. As a result, many questions remain about the cognitive and neural mechanisms of language production and their relationship to language comprehension mechanisms.

Language production encompasses multiple cognitive processes, from formulating a thought to selecting the right words and constructions (lexical access) and putting them together into a well-formed string (phrase-structure building), to retrieving the phonological forms associated with each word, to finally planning and executing a series of motor movements (e.g., articulatory movements for spoken language) (e.g., Levelt, 1989; Goldrick et al., 2014). The latter, lower-level aspects of language production draw on a relatively well-characterized network of superior temporal and frontal areas, including an area in posterior left inferior frontal gyrus (IFG), in line with Broca’s original report (e.g., Hillis et al., 2004; Bohland & Guenther, 2006; Bouchard et al., 2013; Flinker et al., 2015; Fridriksson et al., 2016; Long et al., 2016; Basilakos et al., 2018; Guenther, 2016). This ‘articulation network’ appears to be distinct from what we refer to here as the ‘language network’—the network of (more inferior) temporal and (more anterior) frontal areas that have been implicated in higher-level linguistic processing (cf. Miozzo et al., 2015; Ries et al., 2017; Strijkers et al., 2017 for claims that some articulation areas may contribute to higher-level language production).

The language network can be defined in individual participants using i) functional localizer contrasts (Saxe et al., 2006) like reading sentences vs. nonwords, or listening to speech vs. acoustically degraded speech (e.g., Fedorenko et al., 2010; Scott et al., 2017; Lipkin et al., in press) across diverse languages (Malik-Moraleda, Ayyash et al., 2022), or ii) patterns of inter-voxel functional correlations during naturalistic cognition paradigms (e.g., Braga et al., 2020; see also Blank et al., 2014). These language-responsive brain areas are highly selective for language relative to diverse non-linguistic cognitive processes (e.g., Fedorenko et al., 2011; Monti et al., 2012; Ivanova et al., 2020; see Fedorenko & Varley, 2016 and Fedorenko & Blank, 2020 for reviews) and support both lexical semantic (i.e., relating to the processing of word meanings) and combinatorial processes during language comprehension (e.g., Fedorenko et al., 2010, 2020; Bautista & Wilson, 2016; see e.g., Indefrey & Levelt, 2014 for a meta-analysis).

What role does the language-selective network play in language production? Of course, comprehension and production are intimately linked. Most agree that they draw on the same set of linguistic knowledge representations (e.g., Chomsky, 1965; Pickering & Garrod, 2004). Some further argue that certain computations may be analogous between comprehension and production. For example, Chater et al. (2016; Christiansen & Chater, 2016) have suggested that both rely on chunking—compressing detailed representations in order to pass them down (in production) or up (in comprehension) to the next processing stage. Others argue that the production system is always active during comprehension and serves as a vehicle for linguistic prediction (e.g., Federmeier, 2007; Pickering & Garrod, 2013; Dell & Chang, 2014) and that the comprehension system is always active during production, serving as a monitoring mechanism (e.g., Levelt, 1989; cf. Nozari et al., 2011).

This tight relationship between comprehension and production finds empirical support. For example, syntactic structures that are experienced through listening or reading are more likely to be subsequently produced (e.g., Branigan et al., 2000), and the magnitude of this priming effect is not diminished relative to production-to-production priming (e.g., Tooley & Bock, 2014). Comprehension difficulties tend to arise in structures that are dispreferred during production planning (e.g., Hsiao & MacDonald, 2016). And the same (lexico-semantic and phonological) word knowledge is available at the same early latencies during word production and comprehension (Fairs et al., 2021).

Given this theoretical and empirical backdrop, it seems reasonable to expect that the same mechanisms would support comprehension and production. Indeed, a number of past brain imaging studies have directly compared neural responses during comprehension and production tasks and reported overlap within the language network (e.g., Awad et al., 2007; Menenti et al., 2011; Segaert et al., 2012; Silbert et al., 2014; Giglio et al., 2022; see Indefrey, 2018 and Walenski et al., 2019 for meta-analyses; see Strijkers & Costa, 2016 and Gambi & Pickering, 2017 for theoretical proposals of shared circuitry; cf. Indefrey & Levelt, 2004 and Indefrey, 2011 for arguments for distinct circuits supporting word-level comprehension vs. production). However, these studies and meta-analyses have a) often reported overlap in only some of the language regions, and these regions differ across studies (e.g., Awad et al., 2007 report overlap within anterior temporal cortex and temporo-parietal junction, but Segaert et al., 2012 in posterior temporal and inferior frontal areas), and/or b) have not controlled for lower-level perceptual/motor demands, making the nature of the overlap difficult to interpret (e.g., Silbert et al., 2014). Furthermore, all past studies (cf. Matchin & Wood, 2020; see Discussion) have relied on the traditional random-effects analytic approach where activations are averaged across individuals voxel-wise and inferences are drawn based on the resulting group-level maps. Because of the well-established inter-individual variability in the precise locations of functional areas (e.g., Frost & Goebel, 2012; Tahmasebi et al., 2012)—including language areas (e.g., Fedorenko et al., 2010; Mahowald & Fedorenko, 2016; Braga et al., 2020; Lipkin et al., in press)—and the high functional heterogeneity of the association cortex, where distinct areas often lay adjacent to each other (e.g., Scholz et al., 2009; Fedorenko et al., 2012b; Deen et al., 2015; Braga & Buckner, 2017; Braga et al., 2020; Deen & Freiwald, 2022), this approach can both miss activations and over-estimate activation overlap between nearby areas (e.g., Saxe et al., 2006; Nieto-Castañón & Fedorenko, 2012; Fedorenko, 2021).

Another unresolved question about the architecture of language production concerns the relationship between the mechanisms that support lexical access (word retrieval) vs. those that support phrase-structure building (combining words together to form phrases and sentences). Some past studies that examined neural responses to language production have conflated these two kinds of demands (e.g., Awad et al., 2007; Silbert et al., 2014). Other studies have attempted to separate them and argued for different parts of the language network supporting lexical and combinatorial aspects of language production (e.g., Menenti et al., 2011). However, if a) the same representations subserve comprehension and production, and given that b) for comprehension, lexico-semantic and syntactic processing do not appear to be spatially segregated (e.g., Fedorenko et al., 2020), a dissociation between lexical access and phrase-structure building in production would seem surprising.

Of high relevance to the relationships a) between production and comprehension, and b) between lexical access and combinatorial processing in production is a hypothesis about the existence of mechanisms that selectively support phrase-structure building in production (e.g., Bock, 1982, 1995; Matchin & Hickok, 2019). This hypothesis is motivated by a putative asymmetry between production and comprehension with respect to combinatorial processing. In particular, whereas linearization (putting words in a particular order) and morpho-syntactic agreement processes are necessary aspects of language production (cf. Swets et al., 2013; Goldberg & Ferreira, 2022), comprehension is possible even when word order and other morpho-syntactic cues are degraded or absent (e.g., Ferreira et al., 2002; Ferreira, 2003; Levy, 2008; Levy et al., 2009; Gibson et al., 2013; Ferreira & Lowder, 2016; Mollica et al., 2020; Mahowald, Diachek et al., 2022; cf. Shain et al., in press for evidence that comprehenders compute fine-grained syntactic representations even during passive listening in naturalistic paradigms). Matchin & Hickok (2019), based largely on data from aphasia, recently proposed that such a mechanism may be housed in the inferior frontal component of the language network (in contrast with its posterior temporal component, which they hypothesized supports morpho-syntactic demands in both comprehension and production). However, this proposal has not been thoroughly evaluated to date (cf. the two recent studies that are summarized in the Discussion).

To illuminate the contribution of the language-selective network to language production, we use fMRI to examine the responses of the language areas—defined in individual participants by an extensively validated comprehension-based language localizer (Fedorenko et al., 2010)— during production tasks. To examine both phrase-structure building and lexical access using this *precision fMRI* approach, we adapt a paradigm that has proven fruitful in probing combinatorial and lexico-semantic processes in comprehension (e.g., Friederici et al., 2000; Humphries et al., 2007; Fedorenko et al., 2010, 2012a; Pallier et al., 2011; Shain et al., 2021). In particular, we examine neural responses during spoken (Experiments 1-2) and typed (Experiment 3) production of sentences and lists of words (as well as control, nonword sequences in Experiments 1 and 3). Brain areas that support phrase-structure building should work harder (i.e., exhibit stronger BOLD responses) when words have to be combined (sentence production) compared to retrieval of unrelated words (word-list production), given that sentence production requires additional operations, including semantic composition, ordering the words, and implementing the relevant morpho-syntactic agreement processes. And brain areas that support lexical retrieval should work harder when words must be accessed (word-list production) compared to a simple articulatory task (nonword production) given the additional computations required for word retrieval. (We, of course, acknowledge that both phrase-structure building and lexical access may themselves consist of multiple distinct sub-processes (e.g., some have distinguished between syntactic vs. semantic combinatorial processing; e.g., Hagoort & Indefrey, 2014; Pylkkänen, 2019); here, we probe these aspects of language production using relatively broad contrasts given that this is the first attempt to examine language production using precision fMRI.) To evaluate the hypothesis about brain mechanisms that are selective for phrase-structure building in production relative to comprehension, we additionally included sentence and word-list comprehension conditions.

## Materials and Methods

### Participants

Forty-one individuals (age 18-31, mean 23.3 years; 28 (68.3%) females) from the Cambridge/Boston, MA community participated for payment across three fMRI experiments (n=29 in Experiment 1; n=12 in Experiment 2; and n=14 in Experiment 3; participants in Experiment 3 were a proper subset of the participants in Experiment 1). All were native speakers of English. Of the thirty-two participants for whom handedness data were available, twenty-eight participants (87.5%) were right-handed, as determined by the Edinburgh Handedness Inventory (Oldfield, 1971) or self-report, two (6.25%) were left-handed, and two (6.25%) were ambidextrous (see Willems et al., 2014, for arguments for including non-right-handers in cognitive neuroscience experiments). Handedness data were not recorded for the remaining 9 participants. All but one participant showed typical left-lateralized language activations in the language localizer task; one (right-handed) participant in Experiment 2 showed right-lateralized language activations; we chose to include this participant to err on the conservative side. For Experiment 3, we recruited participants who could type without seeing the written output or the keyboard itself. All participants gave informed written consent in accordance with the requirements of MIT’s Committee on the Use of Humans as Experimental Subjects (COUHES).

### Design, materials, and procedure

Each participant completed a comprehension-based localizer task for the language network (Fedorenko et al., 2010) and a critical language production experiment. All but one participant (in Experiment 1) additionally completed a localizer task for the domain-general Multiple Demand (MD) network (Duncan, 2010, 2013). Because the MD network has been shown to be generally sensitive to task difficulty across domains (e.g., Duncan & Owen, 2000; Fedorenko et al., 2013; Hugdahl et al., 2015; Shashidhara et al., 2019; Assem et al., 2020), activity levels therein can be used to determine the relative difficulty levels of the different production conditions, to aid the interpretation of the results. Some participants also completed one or more tasks for unrelated studies. The scanning sessions lasted approximately two hours.

#### Language network localizer

The regions of the language network were localized using a task described in detail in Fedorenko et al. (2010) and subsequent studies from the Fedorenko lab (and is available for download from https://evlab.mit.edu/funcloc/). Briefly, participants silently read sentences and lists of unconnected, pronounceable nonwords in a blocked design. The sentences > nonwords contrast targets brain regions that that support high-level language comprehension. This contrast generalizes across tasks (e.g., Fedorenko et al., 2010; Scott et al., 2017; Ivanova et al., 2020) and presentation modalities (reading vs. listening; e.g., Fedorenko et al., 2010; Scott et al., 2017; Chen et al., 2021; Malik-Moraleda, Ayyash et al., 2022). All the regions identified by this contrast show sensitivity to lexico-semantic processing (e.g., stronger responses to real words than nonwords) and combinatorial semantic and syntactic processing (e.g., stronger responses to sentences and Jabberwocky sentences than to unstructured word and nonword lists) (e.g., Fedorenko et al., 2010, 2012a, 2016, 2020; Blank et al., 2016; Shain et al., 2021). More recent work further shows that these regions are also sensitive to sub-lexical regularities (Regev et al., 2021), in line with the idea that this system stores our linguistic knowledge, which encompasses regularities across representational grains, from phonological and morphological schemas, to words, to constructions (e.g., Jackendoff, 2002; Jackendoff & Audring, 2019).

Stimuli were presented one word/nonword at a time at the rate of 350-450ms (differing slightly between variants of the localizer; **Table SI-1)** per word/nonword. Participants read the materials passively and performed either a simple button-press or memory probe task at the end of each trial (which were included in order to help participants remain alert). The memory probe required the participant to indicate whether a given word was from the sentence/list of nonwords they had just read. Each participant completed two ~6 min runs. In Experiments 1 and 3, all participants completed the language localizer in the same session as the production experiment. In Experiment 2, 8 participants completed the language localizer in the same session as the production experiment and the remaining 4 participants completed the language localizer in an earlier scanning session (see Mahowald & Fedorenko, 2016 and Braga et al., 2020 for evidence of high across-session reliability of the activation patterns).

#### Multiple Demand (MD) network localizer

The regions of the Multiple Demand (MD) network (Duncan, 2010; Duncan et al., 2020) were localized using a spatial working memory task contrasting a harder condition with an easier condition (e.g., Fedorenko et al., 2011, 2013; Blank et al., 2014). The hard > easy contrast targets brain regions engaged in cognitively demanding tasks. Fedorenko et al. (2013) have established that the regions activated by this task are also activated by a wide range of other demanding tasks (see also Duncan and Owen, 2000; Hugdahl et al., 2015; Shashidhara et al., 2019; Assem et al., 2020).

On each trial (8 s), participants saw a fixation cross for 500 ms, followed by a 3 x 4 grid within which randomly generated locations were sequentially flashed (1 s per flash) two at a time for a total of eight locations (hard condition) or one at a time for a total of four locations (easy condition). Then, participants indicated their memory for these locations in a two-alternative, forced-choice paradigm via a button press (the choices were presented for 1,000 ms, and participants had up to 3 s to respond). Feedback, in the form of a green checkmark (correct responses) or a red cross (incorrect responses), was provided for 250 ms, with fixation presented for the remainder of the trial. Hard and easy conditions were presented in a standard blocked design (4 trials in a 32 s block, 6 blocks per condition per run) with a counterbalanced order across runs. Each run included four blocks of fixation (16 s each) and lasted a total of 448 s. Each participant completed two runs, except for one participant in Experiment 1, who completed one run. In Experiment 1, of the 28 participants who completed the MD localizer, 25 did so in the same session as the production experiment (this included 11 of the 14 participants who also participated in Experiment 3) and the remaining 3 participants completed the MD localizer in an earlier scanning session. In Experiment 2, 9 participants completed the MD localizer in the same session as the production experiment and the remaining 3 participants – in an earlier session.

Like the language localizer, the MD localizer has been extensively validated, and a network that closely corresponds to the one activated by the MD localizer emerges from task-free (resting state) data (e.g., Assem et al., 2020; Braga et al., 2020; also Blank et al., 2014; Malik-Moraleda, Ayyash et al., 2022).

#### General approach for the language production tasks

Tapping mental computations related to high-level language production—including both lexical access and combining words into phrases and sentences—is notoriously challenging because linguistic productions originate from internal conceptual representations (e.g., Levelt, 1989; Bock, 1996; Goldrick et al., 2014). These representations are difficult to probe and manipulate without sacrificing ecological validity. However, given that many open questions remain about how language production is implemented in the mind and brain, and the need for careful comparisons (critical for interpretability; e.g., Mook, 1983), we opted for a controlled experimental approach. In particular, building on a strong foundation of behavioral work on language production, we used pictorial stimuli to elicit object labels and event-level linguistic descriptions.

#### Experiment 1 (Spoken production)

*Design.* Participants were presented with a variety of visual stimuli across six conditions. In the two critical language production conditions—sentence production and word-list production—participants were instructed to speak out loud, but to move their heads as little as possible. The sentence production (SProd) condition is the closest to reflecting the language production demands of everyday life, where we often communicate event-level descriptions using phrases and sentences. In this condition, participants viewed photographs of common events involving humans, animals, and inanimate objects (**Figure 1a-i, 1b-i**) and were asked to produce a description of the event (e.g., “The girl is smelling a flower”). This condition targets sentence-level production planning and execution, which includes a) retrieving the words for the entities/objects and actions, and b) combining them into an utterance, including semantic composition, ordering the words, and implementing the relevant syntactic agreement processes. Of course, participants are not guaranteed to produce complete sentences. In fact, when asked to describe event pictures, participants commonly revert to ‘headlinese’ (the register used in newspaper headlines) and produce descriptions like “girl smelling a flower” or “girl smells flower” (**Figure SI-1**). Importantly, such elliptical productions still require cognitive operations above and beyond lexical retrieval, including semantic composition but also morpho-syntactic planning and execution given that they obey the syntactic constraints of this particular register, including word order and agreement constraints (see e.g., Halliday, 1967; Mårdh, 1980; van Dijk 1988 for discussions of linguistic features of headlinese).

**Figure 1.**
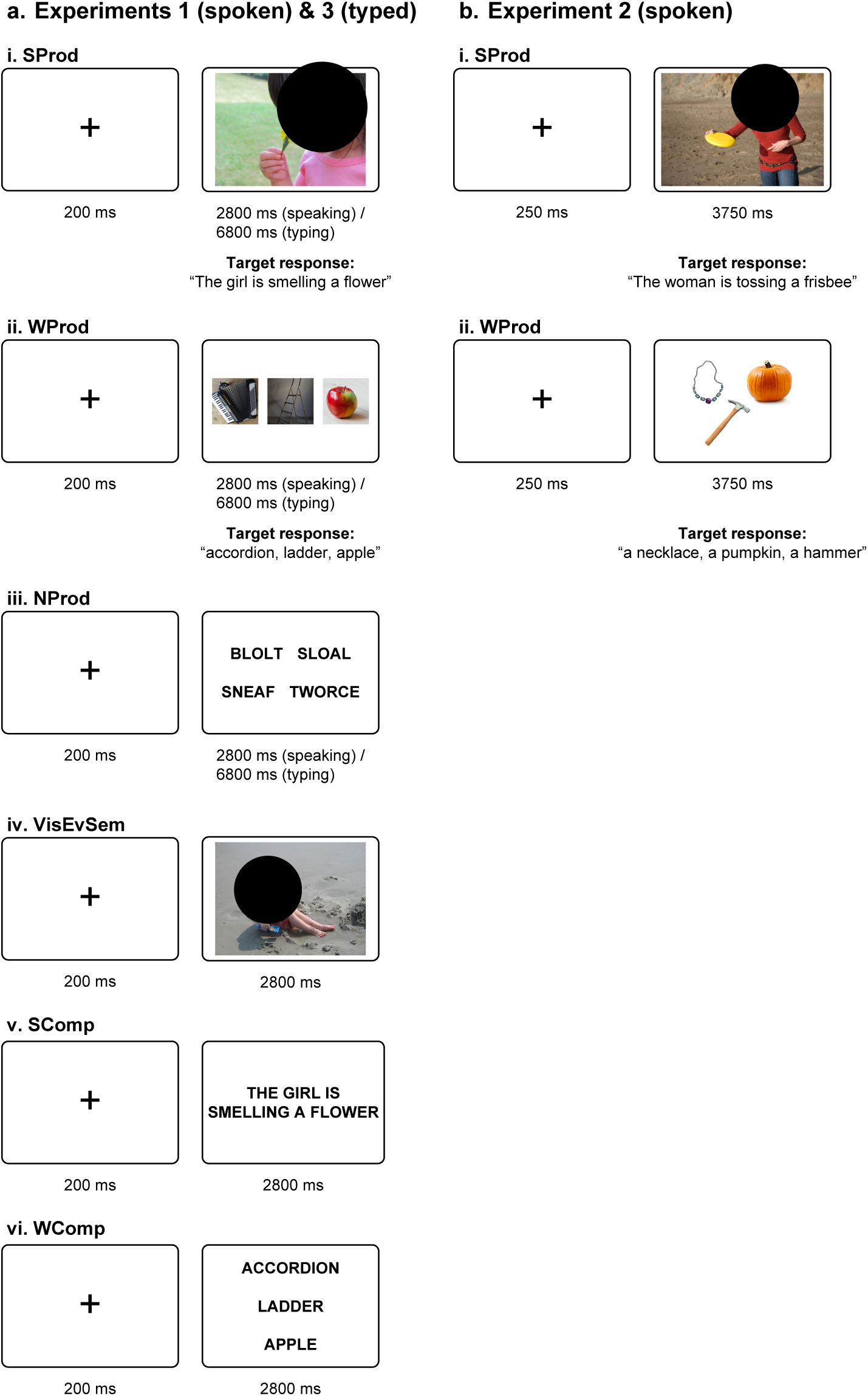
Sample trials for each condition of Experiments 1-3. NOTE: faces have been redacted from event photographs (black circles); for the original images, see https://github.com/jennhu/LanguageProduction/. **a.** In Experiments 1 and 3, participants performed two production tasks: producing descriptions of events depicted in naturalistic photographs (SProd), and producing names of unrelated, isolated objects depicted in separate photographs (WProd). In two control conditions, participants spoke or typed monosyllabic nonwords (NProd), and indicated whether events depicted in photographs took place indoors or outdoors (VisEvSem). Finally, participants performed two comprehension tasks: silently reading sentences (SComp) and word lists (WComp), mirroring the structure and content of target responses from the production trials (see Methods for details). For the production conditions, trials lasted 2800 ms in Experiment 1 (spoken production) and were increased to 6800 ms in Experiment 3 (typed production). **b.** In Experiment 2, participants performed the sentence and word production conditions (SProd, WProd) with a different set of materials.

In the word-list production (WProd) condition, participants viewed groups of 2, 3, or 4 photographs of inanimate objects (**Figure 1a-ii, 1b-ii**) and were asked to name each object in the set (e.g., “accordion, ladder, apple”). The number of objects in each group (2-4) matched the number of content words in the target productions in the SProd condition. This condition targets word-level production planning and execution. To isolate the mental processes related to single-word production, photographs were manually grouped in a way that minimized semantic associations between the objects, to prevent participants from unintentionally forming phrases/clauses with the retrieved words.

The experiment also included two control conditions: low-level (nonword-list) production and semantic judgments about visual events. In the nonword-list production (NProd) condition, participants viewed lists of 4 monosyllabic nonwords (**Figure 1a-iii**) and were asked to say them out loud (e.g., “blolt, sloal, sneaf, tworce”). The nonwords obeyed the phonotactic constraints of English and were selected to be sufficiently distant from phonologically neighboring words. This condition targets low-level articulatory planning and execution and was included as a more stringent baseline (in addition to the low-level fixation baseline) to which sentence and word production conditions could be compared (see e.g., Bohland & Guenther, 2006; Okada & Hickok, 2006; Flinker et al., 2015 for evidence that this kind of condition robustly activates articulation areas). We acknowledge the difference between this condition and the other production conditions in that this condition involves visually presented linguistic strings (cf. pictures in the SProd and WProd conditions); it is difficult to create minimally different conditions one of which does and the other does not require lexical access while having similar lower-level (e.g., articulation) demands. And in the visual event semantics (VisEvSem) condition, participants viewed photographs of events (as in SProd) and were asked to indicate whether the depicted event takes place indoors or outdoors (a relatively high-level judgment that requires visual event perception and also draws on world knowledge) via a two-choice button box (**Figure 1a-iv**). This condition targets visual and conceptual processing of events and was included to ensure that responses to the SProd condition, which uses these pictorial stimuli, were not due to these cognitive processes (see e.g., Ivanova et al., 2021 and Sueoka et al., 2022 for some evidence of engagement of the language areas in visual event semantics).

Finally, the experiment included two reading comprehension conditions: sentence comprehension and word-list comprehension. In both conditions, participants were instructed to read the stimuli silently (as in the language localizer). In the sentence comprehension (SComp) condition, participants viewed short sentences describing common events (e.g., “The girl is smelling a flower”) and were asked to read them. These sentences described the same events that were depicted in the photographs used in the SProd condition (**Figure 1a-v**); however, as is common in sentence-level experiments, any given participant experienced an event once – either as a sentence (SComp condition) or as a photograph (SProd condition) (elaborated in <u>Procedure</u> below). The SComp condition targets sentence-level comprehension processes, including lexico-semantic and combinatorial (semantic and syntactic) processes. In the word-list comprehension (WComp) condition, participants viewed lists of 2, 3, or 4 object names (**Figure 1a-vi**) and were asked to read them (e.g., “accordion, ladder, apple”). This condition targets word-level comprehension. As in the WProd condition, object names in the WComp condition were grouped in a way that minimized semantic associations. These conditions were included to enable comparisons of responses to content-matched sentences and word lists across production and comprehension as relevant to the question of production-selective mechanisms.

*Materials*. To obtain the event photographs for the SProd and VisEvSem conditions, we first manually selected 400 images clearly depicting everyday events from the Flickr30k dataset (Young et al., 2014). We then ran a norming study on Amazon.com’s Mechanical Turk to identify the stimuli that would elicit the most consistent linguistic descriptions across participants. On each trial, participants viewed a single photograph and were given the instructions “Please provide a one-sentence description of what is happening in the photo.” They were able to type freely in a textbox below the image and could only proceed to the next trial after submitting a non-empty response. We recruited n=30 participants for each of the 400 images, and each participant produced descriptions for 100 images.

To analyze the resulting 12,000 responses, we used the Python spaCy natural language processing library (Honnibal et al., 2020) to parse each production into the subject noun phrase (NP), verb phrase (VP), subject NP head, and VP head. After manually cleaning the parses for consistency, we computed three metrics for each photograph: (1) the number of unique responses in each of the parsed categories, (2) the number of unique lemmas for the single-word parsed categories (subject NP head and VP head), and (3) the standard deviation of the number of tokens per production. We then obtained a ‘linguistic variability’ score by summing these three values for each image and chose the 200 photographs with the lowest scores. Finally, we hand-selected 128 from these 200 to maximally cover a range of objects and actions. These photographs were used in the SProd and VisEvSem conditions, and the associated sentence descriptions (the most frequently used description for each photograph) were used in the SComp condition.

For the WProd and WComp conditions, we wanted to use materials that would be semantically (and lexically) similar to the ones used in the SProd and SComp conditions. As a result, to obtain the object photographs for the WProd condition, we first identified between 2 and 4 words in each of the 128 sentence descriptions that referred to inanimate objects (we avoided animate entities like ‘a man’ or ‘a woman’ because in the setup that we used, with multiple objects presented at once, we wanted to avoid the possibility of participants constructing event-level representations). For example, from the description “A man is playing saxophone in a park” we selected ‘saxophone’, and from the description “A man is sitting on a bench reading the newspaper” we selected ‘bench’ and ‘newspaper’. This resulted in a total of 120 words. Next, we selected images of each object from the THINGS database (Hebart et al., 2019) as well as a repository of license-free stock photographs. In those images, each object is presented on a neutral but naturalistic background, which isolates the object from possibly associated events or concepts. We generated all possible groups of 2-, 3-, and 4-object images, and then took a random sample of 40 2-object, 80 3-object, and 40 4-object groups, as there was an average of 3 content words in our target sentence productions. We then manually selected the final 128 object groups by discarding groups with semantically related objects and ensuring that each object appeared 1-3 times. The associated words (grouped in the same way) were used in the WComp condition. The order of objects and words was randomized within each group during presentation.

Finally, the nonwords for the NProd condition were selected from the ARC Nonword Database (Rastle et al., 2002). We began by selecting all the monomorphemic syllables involving orthographically existing onsets, bodies, and legal bigrams. We then obtained the final set of 256 nonwords by filtering for low numbers of onset and phonological neighbors in order to minimize the likelihood of these nonwords priming real words. These 256 nonwords were then randomly distributed into 64 groups of 4.

All the materials for this experiment and Experiment 2 are available on GitHub (https://github.com/jennhu/LanguageProduction); the sentences, word lists, and nonword lists are also provided in the SI for convenience (Appendix A).

*Procedure.* Following the general practice in the field of sentence processing, the same event or object group did not appear in both a production condition and its corresponding comprehension condition for any given participant (in order to avoid influences on the processing of sentence 1 in condition A from the earlier processing of sentence 1 in condition B). To achieve this, we distributed the materials in the SProd, WProd, SComp, and WComp conditions (i.e., 128 event images, 128 corresponding target sentences, 128 object group images, and 128 corresponding target word lists) into two experimental lists. We assigned a unique number 1-128 to each event and object group, such that event image *x* corresponds to sentence *x*, and object group *x* corresponds to word list *x*. These numbers were assigned such that sets 1-64 and 65-128 were each semantically diverse (e.g., two images of a person playing a musical instrument were assigned to different sets). Furthermore, the 2-, 3-, and 4-object groups were evenly distributed across the two sets (1-64 and 65-128). Finally, this numbering was used to create two lists. In List 1, event images 1-64 in SProd appeared with sentences 65-128 in SComp, and similarly object group images 1-64 in WProd appeared with word lists 65-128 in WComp. And in List 2, event/object group images 65-128 in SProd/WProd appeared with sentences/word lists 1-64 in SComp/WComp. The materials for the NProd condition were identical across lists, and the materials for the VisEvSem condition were the same as the SProd materials in that list.

The materials in each condition (and each list, where relevant) were grouped into 16 blocks of 4 trials each; this was done separately for each participant. (Note that although we had enough materials to yield 16 blocks per condition, we ended up presenting 12 blocks per condition for any given participant because—based on pilot participants—this number of blocks per condition gave us sufficient power to elicit clear between-condition differences.) Each block was preceded by instructions, which told the participants what they would be doing in the trials to come: “Describe the event out loud” for SProd, “Name the objects out loud” for WProd, “Say the nonwords out loud” for NProd, “Inside (=1) or outside (=2)?” for VisEvSem, “Read the sentence silently” for SComp, and “Read the words silently” for WComp. The instructions remained on the screen (in small font in the bottom left corner of the screen) throughout the block to minimize the demands associated with holding onto the instructions and to help participants in case they missed the block-initial instructions screen. Each trial lasted 3 sec and consisted of an initial fixation cross (0.2 sec) and stimulus presentation (2.8 sec). In the SProd and VisEvSem conditions, the stimulus was a single event picture; in the WProd condition, the stimulus was a set of 2-4 object pictures (presented all at once); in the NProd condition, the stimulus was a set of 4 nonwords (presented all at once); in the SComp condition, the stimulus was a sentence (presented all at once); and in the WComp, the stimulus was a set of 2-4 words (presented all at once) (see **Figure 1a**). The block-initial instructions were presented for 2 sec. Thus, each block lasted 14 sec (2 sec instructions and 4 trials 3 sec each).

The total of 72 experimental blocks (12 blocks * 6 conditions) were distributed into 6 sets, corresponding to runs, of 12 blocks each (2 blocks per condition). Each run additionally included 3 fixation blocks of 12 sec each: one at the beginning of the run, one after the first six experimental blocks, and one at the end. Thus, each run consisted of 12 experimental blocks of 14 sec each and 3 fixation blocks of 12 sec each, lasting a total of 204 sec (3 min 24 sec). Each participant completed 4-6 runs (for a total of 8-12 blocks per condition). The order of conditions was palindromic within each run and varied across runs and participants.

Prior to entering the scanner, participants were provided with printed instructions and were guided through sample items that mimicked the experimental stimuli. The experimental script with all the materials is available at GitHub: https://github.com/jennhu/LanguageProduction.

#### Experiment 2 (Spoken production; conceptual replication of Experiment 1)

*Design.* Experiment 2 was designed and conducted prior to Experiment 1 and constituted an early attempt (ca. 2012) to develop a word and sentence production paradigm for fMRI. For this reason, it doesn’t include all the critical control conditions that are included in Experiment 1. Nevertheless, it serves as a conceptual replication of the findings from the critical production conditions (SProd and WProd) in Experiment 1 while generalizing the results to a new set of materials and an independent group of participants. The design was identical except that in the word-list production (WProd) condition, participants always viewed groups of 3 object photographs (cf. 2, 3, or 4 object photographs in Experiment 1), and they were asked to name each object in the set with an indefinite article (e.g., “a necklace, a pumpkin, a hammer”), which includes some basic phrase-level combinatorial processing in addition to lexical retrieval. The experiment also included two other conditions that are not directly relevant to the current investigation and are therefore not discussed.

*Materials.* The materials were selected from the publicly available images in the Google Images database and consisted of 96 event images for the SProd condition (**Figure 1b-i**), and 288 object images for the WProd (**Figure 1b-ii**) condition. The event photographs were similar in style to those used in Experiments 1, but were more semantically diverse, including not only humans interacting with inanimate objects (as most events in Experiments 1), but also humans interacting with other humans, and humans interacting with animals. The object photographs were also similar in style to those used in Experiment 1, but did not include any background, and were also more semantically diverse, including not only inanimate objects, but also humans (where the occupation of the person is clear: e.g., a chef, a juggler, a ballerina, etc.) and animals. As in Experiment 1, the object photographs were grouped in a way that minimized semantic associations between the objects.

*Procedure.* The materials in each condition were grouped into 24 blocks of 4 trials each; this was done separately for each participant. (The materials were further divided into two experimental lists of 12 blocks per condition.) Each block was preceded by instructions, which told the participants what they would be doing in the trials to come: “Describe the events” for SProd, and “Name the objects” for WProd. Each trial lasted 4 sec and consisted of an initial fixation cross (0.25 sec) and stimulus presentation (3.75 sec). In the SProd condition, a trial consisted of a single event picture, and in the WProd condition, a trial consisted of three object pictures (presented all at once in a triangular configuration) (see **Figure 1b**). Participants were instructed to describe the events with complete sentences (e.g., “The woman is tossing a frisbee”) and to name the objects with indefinite determiners (e.g., “a necklace, a pumpkin, a hammer”). The block-initial instructions were presented for 2 sec. Thus, each block lasted 18 sec (2 sec instructions and 4 trials 4 sec each).

The total of 48 experimental blocks in each list (12 blocks * 4 conditions, 2 of which are of interest to the current study) were distributed into 4 sets, corresponding to runs, of 12 blocks each (3 blocks per condition). Each run additionally included 4 fixation blocks of 18 sec each: one at the beginning of the run, and one after each set of four experimental blocks. Thus, each run consisted of 12 experimental blocks of 18 sec each and 4 fixation blocks of 18 sec each, lasting a total of 288 sec (4 min 48 sec). Eleven participants completed 4 runs (for a total of 12 blocks per condition) and one participant completed 2 runs (for a total of 6 blocks per condition). The order of conditions was palindromic within each run and varied across runs and participants. Prior to entering the scanner, participants were provided with printed instructions and were guided through sample items that mimicked the experimental stimuli. The experimental script with all the materials is available at GitHub: https://github.com/jennhu/LanguageProduction.

#### Experiment 3 (Typed production; extension to another output modality)

*Design and materials.* Experiment 3 served to generalize the results from Experiment 1 to another output modality and was performed by 14 of the 29 participants in Experiment 1. If the data patterns observed for spoken production reflect cognitive processes related to high-level aspects of language production (i.e., lexical access and phrase-structure building), then they should be similar regardless of whether the utterances are spoken or written. This logic is similar to the logic in past studies of language comprehension that have compared neural responses to spoken and written (or signed for sign languages) linguistic input. Such studies have found that the high-level frontal and temporal language areas are sensitive to lexico-semantic and combinatorial processing during comprehension, regardless of the input modality (e.g., Fedorenko et al., 2010, 2016; Vagharchakian et al., 2012; Regev et al., 2013; Scott et al., 2017; Deniz et al., 2019). The design of Experiment 3 was identical to that of Experiment 1, except that in the production conditions (the two critical conditions—SProd and WProd— and the NProd control condition), participants were asked to type their responses on a scanner-safe keyboard (described below) instead of saying them out loud. For the control VisEvSem condition, participants were asked to type their answers (1 or 2) on the keyboard instead of the button box. The two critical production conditions target the same cognitive processes as in Experiment 1, and the control NProd condition targets low-level hand motor planning and execution. Given that the participants in this experiment also participated in Experiment 1, for any given participant, a different experimental list was used than the list used for Experiment 1 (see Experiment 1 for details of how the lists were constructed).

*Procedure.* The procedure for Experiment 3 only differed from that of Experiment 1 in the trial timing for the three production conditions (SProd, WProd, and NProd). In particular, for these conditions, trial duration was increased from 3 to 7 sec (0.2 sec fixation and 6.8 stimulus presentation) given that typing takes longer than speaking (especially when typing in an unusual position, as described below). Each run therefore consisted of 12 experimental blocks (6 were 14 sec each, as in Experiment 1, and 6 were 23 sec each) and 3 fixation blocks of 12 sec each, lasting a total of 300 seconds (5 min). The on-screen instructions for the production conditions were also adjusted to reflect the difference in output modality: “Type a description of the event” for SProd, “Type the names of the objects” for WProd, and “Copy the nonwords (typing)” for NProd. Each participant completed 6 runs (for a total of 12 blocks per condition). The order of experiments (Experiment 1 and Experiment 3) was counterbalanced across participants.

To collect the typed responses, we built a custom MR-safe wireless keyboard. We purchased an off-the shelf wireless keyboard (Inland model ic210) and removed all the ferrous mechanical parts, such as the case screws and the steel wires used to stabilize the wide keys (Shift, Return, and space keys). We then replaced the highly ferrous alkaline AA battery and pulse width modulated step-up voltage regulator with a lithium ion polymer (LiPo) battery and a linear low-drop out voltage regulator. The keyboard uses silicon dome switches and flexible conductive traces that were not found to be ferrous. The wireless USB receiver was plugged into the MRI suite’s penetration panel through a USB to DB9 filter to prevent the introduction of RF interference into the MR images. The absence of RF interference introduced by the keyboard was confirmed by collecting time series of BOLD scans with and without the presence of the keyboard and keys being pressed during these scans and calculating the pixel-by pixel temporal SNR (tSNR) on a static quality assurance phantom.

During the experiment, the keyboard was placed directly on the participant’s abdomen or on a small non-ferrous platform placed on their abdomen, so they could quite comfortably type while lying in the scanner (akin to working on one’s laptop in bed); however, they were unable to see the output of their typing or their own keystrokes. Participants were given a chance to practice typing prior to the experiment to get accustomed to the setup and the keyboard layout. We collected and monitored the participants’ productions on a computer outside the scanning room.

The experimental script with all the materials is available at GitHub: https://github.com/jennhu/LanguageProduction.

## fMRI data acquisition, preprocessing, and first-level modeling

### Data acquisition

Whole-brain structural and functional data were collected on a whole-body 3 Tesla Siemens Trio scanner with a 32-channel head coil at the Athinoula A. Martinos Imaging Center at the McGovern Institute for Brain Research at MIT. T1-weighted structural images were collected in 176 axial slices with 1 mm isotropic voxels (repetition time (TR) = 2,530 ms; echo time (TE) = 3.48 ms). Functional, blood oxygenation level-dependent (BOLD) data were acquired using an EPI sequence with a 90° flip angle and using GRAPPA with an acceleration factor of 2; the following parameters were used: thirty-one 4.4 mm thick near-axial slices acquired in an interleaved order (with 10% distance factor), with an in-plane resolution of 2.1 mm × 2.1 mm, FoV in the phase encoding (A >> P) direction 200 mm and matrix size 96 × 96 voxels, TR = 2,000 ms and TE = 30 ms. The first 10 s of each run were excluded to allow for steady state magnetization.

### Preprocessing

fMRI data were analyzed using SPM12 (release 7487), CONN EvLab module (release 19b), and other custom MATLAB scripts. Each participant’s functional and structural data were converted from DICOM to NIFTI format. All functional scans were coregistered and resampled using B-spline interpolation to the first scan of the first session (Friston et al., 1995). Potential outlier scans were identified from the resulting subject-motion estimates as well as from BOLD signal indicators using default thresholds in CONN preprocessing pipeline (5 standard deviations above the mean in global BOLD signal change, or framewise displacement values above 0.9 mm; Nieto-Castañón, 2020). Functional and structural data were independently normalized into a common space (the Montreal Neurological Institute [MNI] template; IXI549Space) using SPM12 unified segmentation and normalization procedure (Ashburner & Friston, 2005) with a reference functional image computed as the mean functional data after realignment across all timepoints omitting outlier scans. The output data were resampled to a common bounding box between MNI-space coordinates (−90, −126, −72) and (90, 90, 108), using 2mm isotropic voxels and 4th order spline interpolation for the functional data, and 1mm isotropic voxels and trilinear interpolation for the structural data. Last, the functional data were smoothed spatially using spatial convolution with a 4 mm FWHM Gaussian kernel.

### First-level modeling

For both the language localizer task and the critical experiments, effects were estimated using a General Linear Model (GLM) in which each experimental condition was modeled with a boxcar function convolved with the canonical hemodynamic response function (HRF) (fixation was modeled implicitly, such that all timepoints that did not correspond to one of the conditions were assumed to correspond to a fixation period). Temporal autocorrelations in the BOLD signal timeseries were accounted for by a combination of high-pass filtering with a 128 seconds cutoff, and whitening using an AR(0.2) model (first-order autoregressive model linearized around the coefficient a=0.2) to approximate the observed covariance of the functional data in the context of Restricted Maximum Likelihood estimation (REML). In addition to experimental condition effects, the GLM design included first-order temporal derivatives for each condition (included to model variability in the HRF delays), as well as nuisance regressors to control for the effect of slow linear drifts, subject-motion parameters, and potential outlier scans on the BOLD signal.

### Definition of the language network functional regions of interest (fROIs)

For each participant, we defined a set of language fROIs using group-constrained, subject-specific localization (Fedorenko et al., 2010). In particular, each individual map for the *sentences > nonwords* contrast from the language localizer was intersected with a set of six binary masks. These masks (**Figure 2a**; available at https://evlab.mit.edu/funcloc/) were derived from a probabilistic activation overlap map for the same contrast in a large set of participants (n=220) using watershed parcellation, as described in Fedorenko et al. (2010) for a smaller set of participants. These masks covered the fronto-temporal language network in the left hemisphere. Within each mask, a participant-specific language fROI was defined as the top 10% of voxels with the highest *t*-values for the localizer contrast. (Note that we here included the language fROI within the angular gyrus (AngG) even though this fROI consistently patterns differently from the rest of the language fROIs in its functional response profile and patterns of functional correlations. In other recent papers, we have started excluding the AngG language fROI to focus on the core set of five language fROIs. However, we chose to include it here given the importance of the ‘dorsal stream’—white matter tracts of the arcuate and/or superior longitudinal fasciculus that connect posterior-most temporal/parietal language areas and inferior frontal language areas—in language production (e.g., Hickok & Poeppel, 2004; Fridriksson et al., 2016).)

**Figure 2.**
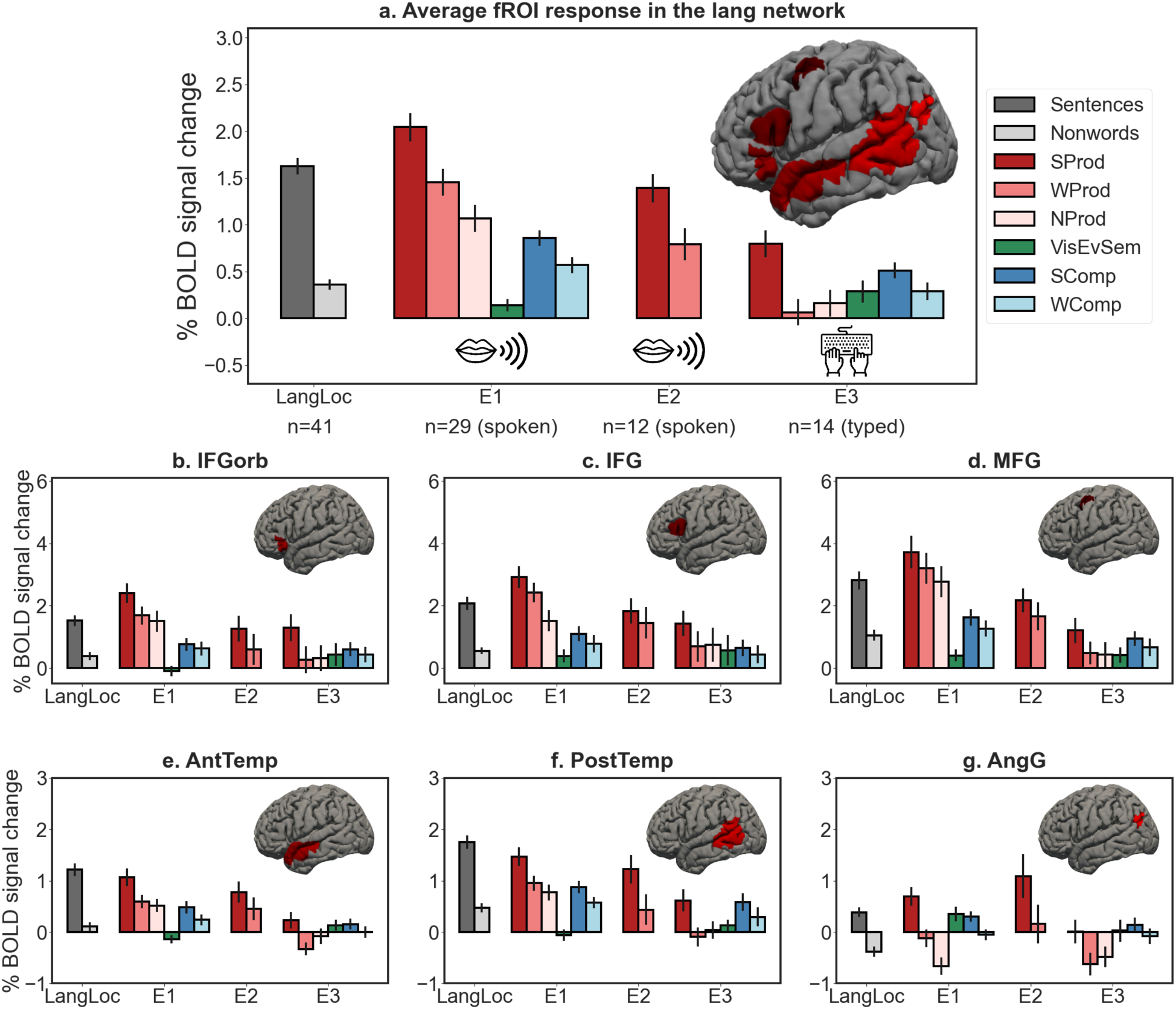
Responses in the language network. Responses in the language network to the language localizer conditions (dark grey=sentences, light grey=nonwords; the responses are pooled across all participants) and the conditions of the critical experiments (red shades=production conditions (from darker to lighter: sentence production (SProd), word-list production (WProd), and nonword-list production (NProd); green=visual event semantic condition (VisEvSem); blue shades=comprehension conditions (dark=sentence comprehension (SComp), light=word-list comprehension (WComp)) in Experiments 1-3. The top panel shows the responses averaging across the six regions of the language network. On the brain inset, we show the parcels that were used to define the individual language functional ROIs (any individual fROI is 10% of the parcel, as described in <u>Methods</u>). The bottom panels show the responses for each of the six regions of the language network. Error bars represent standard errors of the mean over participants.

### Definition of the MD network fROIs

For each participant (except for one participant in Experiment 1 who did not perform the relevant localizer), we defined a set of MD fROIs using group-constrained, subject-specific localization (Fedorenko et al., 2010). Each individual map for the *hard > easy spatial working memory* contrast from the MD localizer was intersected with a set of twenty binary masks (10 in each hemisphere). These masks (**Figure 3a**; available at https://evlab.mit.edu/funcloc/) were derived from a probabilistic activation overlap map for the same contrast in a large set of participants (n=197) using watershed parcellation. The masks covered the frontal and parietal components of the MD network (Duncan, 2010, 2013) bilaterally. Within each mask, a participant-specific MD fROI was defined as the top 10% of voxels with the highest *t*-values for the localizer contrast.

**Figure 3.**
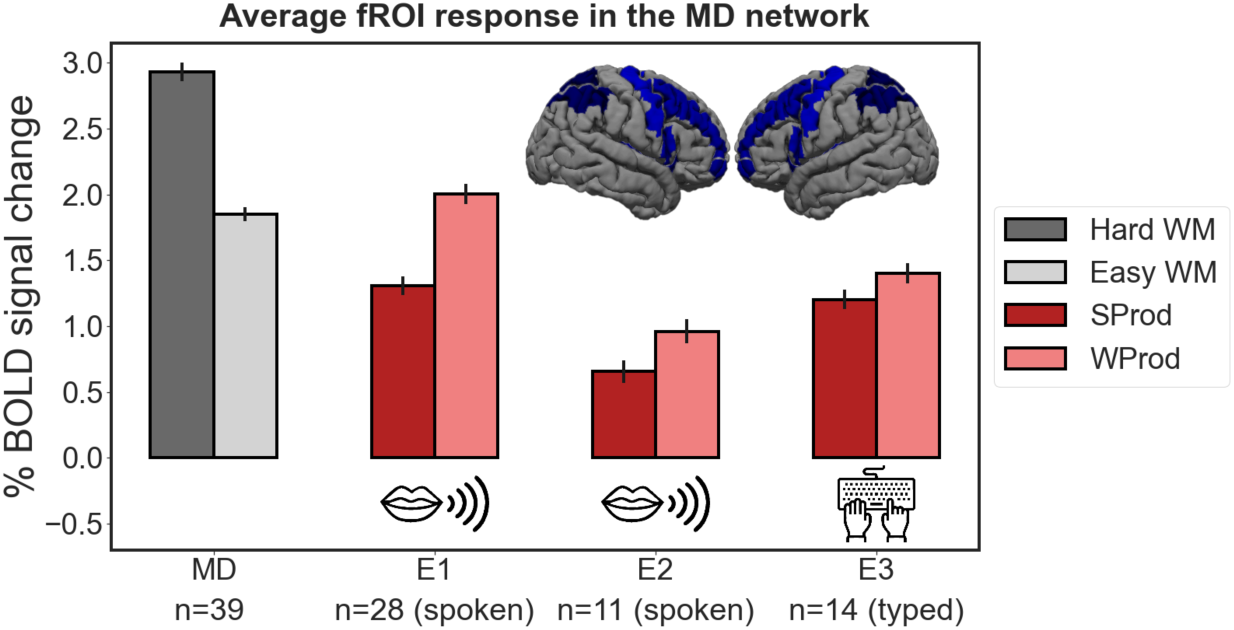
Responses in the MD network. Responses in the bilateral multiple demand (MD) network to the MD localizer conditions (dark grey=hard spatial working memory (Hard WM), light grey=easy spatial working memory (Easy WM); the responses are pooled across all participants) and the production conditions of the critical experiments (from darker to lighter red shades: sentence production (SProd), word-list production (WProd), and nonword-list production (NProd)) in Experiments 1-3. Responses are averaged across the 20 regions of the MD network. On the brain inset, we show the parcels that were used to define the individual MD functional ROIs (any individual fROI is 10% of the parcel, as described in <u>Methods</u>). Error bars represent standard errors of the mean over participants.

## Analyses

All analyses were performed with linear mixed-effects models using the “lme4” package in R (version 1.1.26; Bates et al., 2015) with *p*-value approximation performed by the “lmerTest” package (version 3.1.3; Kuznetsova et al., 2017) and effect sizes (Cohen’s *d*) estimated by the “EMAtools” package (version 0.1.3; Kleiman, 2017).

### Validation of the language and MD fROIs

To ensure that the language and MD fROIs behave as expected (i.e., show a reliable localizer contrast effect), we used an across-runs cross-validation procedure (e.g., Nieto-Castañón & Fedorenko, 2012). In these analyses, the first run of the localizer was used to define the fROIs, and the second run to estimate the responses (in percent BOLD signal change, PSC) to the localizer conditions, ensuring independence (e.g., Kriegeskorte et al., 2009); then the second run was used to define the fROIs, and the first run to estimate the responses; finally, the extracted magnitudes were averaged across the two runs to derive a single response magnitude for each of the localizer conditions. Statistical analyses were performed on these extracted PSC values. As expected, the language fROIs showed a robust *sentences > nonwords* effect (*p*s<10^-8^, |*d|*s>2.05; *p*-values corrected for the number of fROIs using the False Discovery Rate (FDR) correction; Benjamini & Yekutieli, 2001), and the MD fROIs showed a robust *hard > easy spatial working memory* effect (*p*s<10^-6^, |*d|*s>1.84). For participants with a single run of the MD localizer, the activation maps were visually inspected to ensure they look as expected.

### Critical analyses

To estimate the responses in the language fROIs (and MD fROIs for one analysis) to the conditions of the critical experiments, the data from all the runs of the language (or MD) localizer were used to define the fROIs, and the responses to each condition were then estimated in these regions by averaging the effects across the voxels that comprise each individual fROI. Statistical analyses were performed on these PSC values.

For each relevant contrast (as described in detail below, in 1-3), we used two types of linear mixed-effect regression models: i) the network-wise model, which examined the language network as a whole; and ii) the fROI-wise models, which examined each language fROI separately, to paint a more detailed picture. As discussed in the Introduction, treating the language network as an integrated system is reasonable given that the regions of this network a) show similar functional profiles, both with respect to their selectivity for language (e.g., Fedorenko et al., 2011; Fedorenko & Blank, 2020) and their role in lexico-semantic and combinatorial processing during language comprehension (e.g., Fedorenko et al., 2010, 2012, 2016, 2020; Bautista & Wilson, 2016; Blank et al., 2016); and b) exhibit strong inter-region correlations in their activity during naturalistic cognition paradigms (e.g., Blank et al., 2014; Paunov et al., 2019; Braga et al., 2020) and in key functional markers, like the strength of response or the extent of activation in response to language stimuli (e.g., Mahowald & Fedorenko, 2016; Mineroff, Blank et al., 2018). However, to examine potential differences among the language regions with respect to their role in language production, we supplement the network-wise analyses with the analyses of the six language fROIs separately. For the analyses of the MD network, we only use network-wise models given that we use the MD network for control purposes and do not have any hypotheses about differences among the MD regions with respect to the current experiments.

For the network-wise analyses, we fit a linear mixed-effect regression model, predicting the level of BOLD response in the language (or MD) fROIs in the contrasted conditions (as detailed in 1-3 below). The model included a fixed effect for condition and random intercepts for fROI and participant.

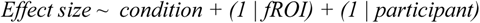

For the fROI-wise analyses, we fit a linear mixed-effect regression model, predicting the level of BOLD response in the target fROI in the contrasted conditions. The model included a fixed effect for condition and a random intercept of participant. The results were FDR-corrected for the number of fROIs.

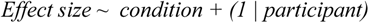

To characterize the responses in the language network to language production, we asked the following three questions.

### 1. Does the language network respond to sentence production?

First, we asked whether sentence production elicits a response in the language network. As discussed in the Introduction, overlap between the language comprehension system (targeted by our language localizer; Fedorenko et al., 2010) and the language production system is expected based on a) current theorizing about their relationship, b) evidence from psycholinguistic and neurophysiological studies, and c) previous PET/fMRI reports of overlap in some of the language regions in studies that have relied on the group-averaging approach (e.g., Awad et al., 2007; Menenti et al., 2011; Segaert et al., 2012; Silbert et al., 2014; Giglio et al., 2022). We also asked whether this response is ubiquitous across the language network or restricted to some of its regions, as some past studies and meta-analyses have suggested (e.g., Awad et al., 2017; Segaert et al., 2012; Giglio et al., 2022; Walenski et al., 2019), and whether it generalizes across output modalities.

To evaluate this question, we examined four contrasts. First, we compared the responses to the sentence production condition (SProd in each experiment) against the fixation baseline. (Although fixation is a liberal baseline, it is important to note that many perceptual and cognitive conditions do not elicit a response in the language network above the fixation baseline, including executive tasks (Blank & Fedorenko, 2020) and music perception (Chen et al., 2021).) Second, we compared the responses to the sentence production condition against the response to the nonword strings condition from the language localizer—an unstructured and meaningless linguistic stimulus. Third and fourth, in Experiments 1 and 3, we compared the responses to the sentence production condition against the response to the low-level production condition (NProd) and the visual event semantics condition (VisEvSem). A brain region that supports sentence production should exhibit a response during the SProd condition (across output modalities) that falls above both the fixation baseline, the nonword strings condition from the language localizer, and the low-level production condition. And if that brain region responds to language production demands rather than the visual and/or semantic processing associated with the processing of event pictures (cf. Ivanova et al., 2021, in prep.), it should also respond more strongly during sentence production than during a semantic task on the same pictures.

### 2. Does the language network contribute to phrase-structure building, lexical access, or both?

Focusing primarily on the well-powered Experiment 1, we probed the responses in the language network to two core aspects of high-level language production. As discussed in the Introduction, if the knowledge representations are shared between comprehension and production, and given that for comprehension, all regions of the language network respond to both lexico-semantic and combinatorial linguistic demands (e.g., Fedorenko et al., 2020), we might expect a similar picture to hold for production. To test the responses of the language regions to phrase-structure building demands above and beyond those associated with single word retrieval, we compared the responses to the sentence production condition (SProd) against the word-list production condition (WProd). And to test the responses to lexical access demands above and beyond lower-level articulation demands, we compared the responses to the word-list production condition (WProd) against the nonword-list production condition (NProd). A brain region that contributes to phrase-structure building should show a SProd>WProd effect, and a brain region that contributes to lexical access should show a WProd>NProd effect.

In a control analysis, we asked whether the SProd condition might elicit stronger responses than the WProd condition because it is more cognitively demanding. To test this possibility, we examined the responses to these conditions in a set of brain regions that have been previously established to be robustly sensitive to general cognitive effort across domains: the regions of the fronto-parietal Multiple Demand (MD) network (Duncan, 2010, 2013; Fedorenko et al., 2013; Hudgdahl et al., 2015; Shashidhara et al., 2019; Assem et al., 2020). This network is functionally distinct from the language network (see Fedorenko & Blank, 2020 for review), and appears to respond during linguistic tasks only in the presence of external task demands, at least for language comprehension (Diachek, Blank, Siegelman et al., 2020; Fedorenko & Shain, 2021).

### 3. Do any brain regions selectively support phrase-structure building during language production relative to comprehension?

Finally, we evaluated the idea from the literature about an asymmetry between production and comprehension: namely, that morpho-syntactic aspects of phrase-structure building are relatively more important for production than comprehension (e.g., Bock, 1982, 1995; Matchin & Hickok, 2019). For example, Bock (1982) writes, “many of the aspects of a sentence’s surface form appear to play a relatively minor role in comprehension”; sentence production, on the other hand, “requires the paraphernalia of the correct morphology, constituent structure and order”. This hypothesis postulates the existence of some production-selective mechanisms. Matchin & Hickok (2019) advocate a version of this idea where the inferior frontal component of the language network selectively/preferentially supports morpho-syntactic demands in production (what they call “morpho-syntactic linearization”, or transforming “nonsequential conceptual information” into “sequences of morphemes”), in contrast to the posterior temporal component, which is hypothesized to support morpho-syntactic demands (“hierarchical structuring”) in both comprehension and production. Although the SProd>WProd contrast in the current study does not *isolate* morpho-syntactic aspects of combinatorial linguistic processing (i.e., it also includes semantic compositional aspects), it certainly *includes* those aspects. To evaluate the prediction of Matchin & Hickok’s proposal, we examined the responses of the IFG and PostTemp language fROIs (and other language fROIs, for completeness) to the SProd, WProd, SComp, and WComp conditions. The sizes of the SComp>WComp and SProd>WProd effects are predicted to be similar for the PostTemp fROI, and the SProd>WProd effect is predicted to be larger than the SComp>WComp effect in the IFG fROI, as would be evidenced by an interaction between task (production vs. comprehension) and stimulus (sentences vs. word lists). (If such an interaction were to obtain in the IFG fROI, a claim about a difference in the profiles of the IFG and the PostTemp fROIs would further require a three-way interaction between task, stimulus, and fROI.)

In addition, we evaluated the production/comprehension asymmetry idea more broadly by asking whether any brain regions—including outside the core left-hemisphere language network—show selective responses to computations related to phrase-structure building during production relative to comprehension. To do so, we searched across the brain for regions that respond more strongly during the SProd condition than each of the WProd and SComp conditions. The SProd>WProd contrast targets voxels that are sensitive to phrase-structure building demands during language production—the aspect of production that is hypothesized to engage selective mechanisms, and the SProd>SComp contrast targets voxels that respond more strongly during sentence production than during sentence comprehension. For this search, we used a whole-brain group-constrained, subject-specific approach (Fedorenko et al., 2010), which is akin to the traditional whole-brain random-effects analysis (Holmes & Friston, 1998), but is more statistically powerful and robust (Blank, 2020; Blank et al., in prep.) given that it a) takes into account inter-individual variability in the precise locations of functional areas, and b) has built into it an across-runs cross-validation procedure to ensure that the regions that emerge show replicable responses over time. Using data from Experiment 1, we created for each participant a whole-brain map that represented a conjunction of contrasts (SProd>WProd and SProd>SComp; for each contrast, we selected the top 10% of most responsive voxels across the brain; the results were similar when selecting voxels based on fixed significance thresholds). Each participant’s map was binarized, with ones corresponding to voxels that fall in the top 10% of voxels for both contrasts above, and zeros otherwise. These individual maps were then overlaid, and watershed parcellation was performed, as described in Fedorenko et al. (2010; see also Julian et al., 2012), to search for areas that that show spatially consistent responses in the majority of participants. The resulting regions were then used as masks to define the individual fROIs using the same two contrasts, selecting the top 10% of voxels based on the *t*-values for each contrast in each parcel and taking the intersection of those voxel sets (the n% approach allows for the definition of the fROIs in each individual). Finally, across-runs cross-validation was used to estimate the responses to the critical conditions (SProd, WProd, and SComp) in these individually defined fROIs and to test for the replicability of the SProd>WProd and SProd>SComp contrasts. The responses of the fROIs with replicable response profiles were then examined with respect to the full range of experimental conditions (including the localizer conditions) to evaluate whether any region(s) is/are indeed selective for phrase-structure building during language production.

## Results

### 1. Every region of the language network responds robustly to sentence production (across output modalities)

In line with past studies (e.g., Awad et al., 2007; Menenti et al., 2011; Siegart et al., 2012; Silbert et al., 2014; Giglio et al., 2022), *spoken sentence production* elicited a robust response in the language network across the two spoken production experiments. This response was stronger than i) the fixation baseline (Experiment 1: d=1.786, p<0.001; Experiment 2: d=1.731, p<0.001), ii) the nonword reading control condition from the language localizer (Experiment 1: d=1.384, p<0.001; Experiment 2: d=1.587, p<0.001), iii) the low-level production condition (Experiment 1: d=0.797, p<0.001), and iv) the visual event semantic processing condition (Experiment 1: d=1.514, p<0.001) (**Figure 2a, Table 1, Table SI-2**). (These effects held in all individual language fROIs (**Figure 2b-g, Table 1, Table SI-2**).)

**Table 1.**
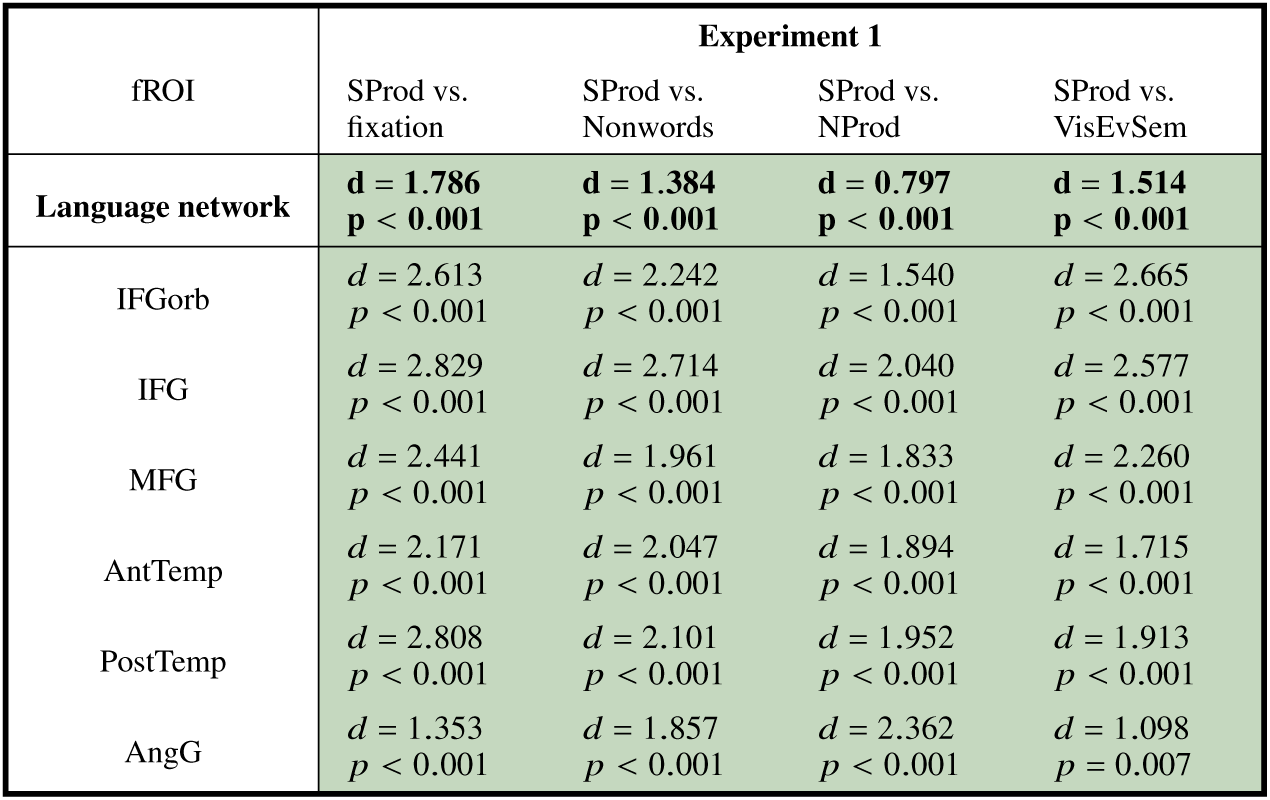
Responses in the language network to spoken sentence production. Effect sizes (Cohen’s d) and estimated p-values for the effect of the spoken SProd condition (relative to four baselines) in linear mixed-effects regression models in Experiment 1 (see Analyses, Q1). Models were fit to perform pairwise comparisons of sentence production (SProd) vs. each of the following conditions: fixation, nonword comprehension (Nonwords; from the language localizer), nonword production (NProd), and visual event semantics processing (VisEvSem). The results are shown averaged across the language network (top row), as well as at the level of individual functional ROIs in the language network (bottom six rows; FDR corrected). Green cells highlight significance at p<0.05 in the predicted direction.

| fROI | Experiment 1 |  |  |  |
| --- | --- | --- | --- | --- |
|  | SProd vs. fixation | SProd vs. Nonwords | SProd vs. NProd | SProd vs. VisEvSem |
| **Language network** | **$d = 1.786$**<br>**$p < 0.001$** | **$d = 1.384$**<br>**$p < 0.001$** | **$d = 0.797$**<br>**$p < 0.001$** | **$d = 1.514$**<br>**$p < 0.001$** |
| IFGorb | $d = 2.613$<br>$p < 0.001$ | $d = 2.242$<br>$p < 0.001$ | $d = 1.540$<br>$p < 0.001$ | $d = 2.665$<br>$p < 0.001$ |
| IFG | $d = 2.829$<br>$p < 0.001$ | $d = 2.714$<br>$p < 0.001$ | $d = 2.040$<br>$p < 0.001$ | $d = 2.577$<br>$p < 0.001$ |
| MFG | $d = 2.441$<br>$p < 0.001$ | $d = 1.961$<br>$p < 0.001$ | $d = 1.833$<br>$p < 0.001$ | $d = 2.260$<br>$p < 0.001$ |
| AntTemp | $d = 2.171$<br>$p < 0.001$ | $d = 2.047$<br>$p < 0.001$ | $d = 1.894$<br>$p < 0.001$ | $d = 1.715$<br>$p < 0.001$ |
| PostTemp | $d = 2.808$<br>$p < 0.001$ | $d = 2.101$<br>$p < 0.001$ | $d = 1.952$<br>$p < 0.001$ | $d = 1.913$<br>$p < 0.001$ |
| AngG | $d = 1.353$<br>$p < 0.001$ | $d = 1.857$<br>$p < 0.001$ | $d = 2.362$<br>$p < 0.001$ | $d = 1.098$<br>$p = 0.007$ |

Moreover, the effects generalized to the typed output modality. *Typed sentence production* (Experiment 3) also elicited a stronger response in the language network than i) fixation (d=1.027, p<0.001), ii) nonword reading (d=0.498, p=0.003), iii) low-level production (d=0.705, p<0.001), and iv) visual event semantic processing (d=0.576, p<0.001) (**Figure 2a, Table SI-2**).

### 2. The language network contributes to both phrase-structure building and lexical access, but phrase-structure building elicits a stronger and more spatially extensive response

The language network responded to both phrase-structure building and lexical access. At the network level, sentence production (SProd) elicited a stronger response than word-list production (WProd) (d=0.499, p<0.001; **Figure 2, Table 2**). This effect replicated in Experiment 2 (d=0.609, p<0.001) and Experiment 3 (d=0.830, p<0.001) (**Figure 2, Table SI-3**). Further, in Experiment 1, this effect was reliable in each of the six fROIs (**Figure 2, Table 2**). Similarly, at the network level, word-list production (WProd) elicited a stronger response than nonword production (NProd) (d=0.325, p=0.004) (**Figure 2, Table 2**; see **Figure SI-2** for a control analysis that shows that the NProd condition elicits the expected strong responses in the premotor and motor cortex), but unlike the SProd>WProd effect, the WProd>NProd effect was only reliable in three of the six fROIs. Further, at the network level and in five of the six fROIs (the IFG language fROI being the exception), the SProd>WProd effect was numerically larger than the WProd>NProd effect, which suggests that the response to sentence production is more strongly driven by phrase-structure building demands. (Interestingly, the region where lexical access demands elicited a larger effect than phrase-structure building demands was the IFG language fROI—a region that has been associated with *combinatorial*, not lexical, processing in much prior literature (e.g., Friederici 2002, 2011; Hagoort, 2005).)

**Table 2.**
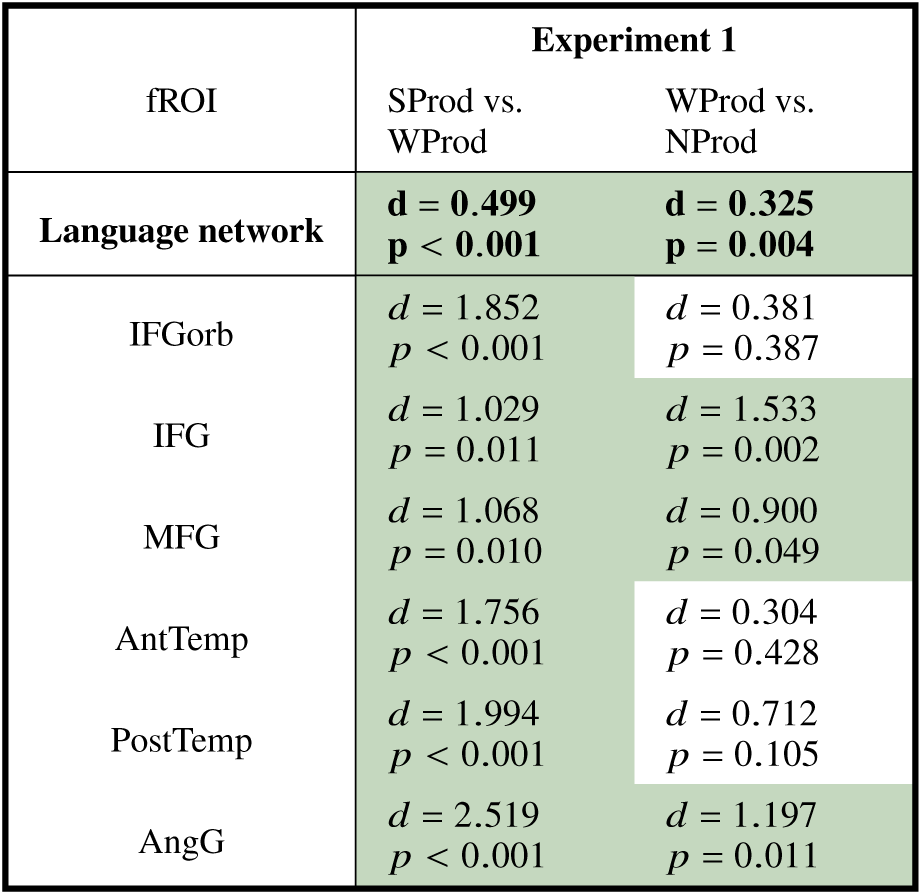
Responses in the language network to phrase-structure building and lexical access. Effect sizes (Cohen’s d) and estimated p-values for the effects associated with phrase-structure building and lexical access in linear mixed-effects regression models in Experiment 1 (see Analyses, Q2). Models were fit to perform pairwise comparisons of sentence production (SProd) vs. word-list production (WProd), and word-list production vs. nonword production (NProd). The results are shown averaged across the language network (top row), as well as at the level of individual functional ROIs in the language network (bottom six rows; FDR corrected). Green cells highlight significance at p<0.05 in the predicted direction.

| fROI | Experiment 1 |  |
| --- | --- | --- |
|  | SProd vs.<br>WProd | WProd vs.<br>NProd |
| **Language network** | **$d = 0.499$**<br>**$p < 0.001$** | **$d = 0.325$**<br>**$p = 0.004$** |
| IFGorb | $d = 1.852$<br>$p < 0.001$ | $d = 0.381$<br>$p = 0.387$ |
| IFG | $d = 1.029$<br>$p = 0.011$ | $d = 1.533$<br>$p = 0.002$ |
| MFG | $d = 1.068$<br>$p = 0.010$ | $d = 0.900$<br>$p = 0.049$ |
| AntTemp | $d = 1.756$<br>$p < 0.001$ | $d = 0.304$<br>$p = 0.428$ |
| PostTemp | $d = 1.994$<br>$p < 0.001$ | $d = 0.712$<br>$p = 0.105$ |
| AngG | $d = 2.519$<br>$p < 0.001$ | $d = 1.197$<br>$p = 0.011$ |

In contrast to the language network, the Multiple Demand network responded more strongly to the WProd condition than the SProd condition (d=0.692, p<0.001), providing evidence that word-list production is, in fact, more cognitively demanding than sentence production, and ruling out general cognitive difficulty as the explanation of the SProd>WProd effect in the language network (**Figure 3**). This effect replicated in Experiment 2 (d=0.353, p<0.001) and Experiment 3 (d=0.225, p=0.01). It is worth noting that strong responses in the MD network during single-word production underscore the contributions of domain-general executive mechanisms to performance in confrontation naming tasks—one of the most commonly used clinical language assessment tools (e.g., Kaplan et al., 1983).

### 3. No evidence of brain regions that selectively support phrase-structure building during language production relative to comprehension

We do not find support for Matchin & Hickok’s (2019) proposal about the IFG language fROI selectively supporting phrase-structure building during production relative to comprehension. As shown in **Figure 4a-b**, contra this proposal’s prediction (illustrated in **Figure 4c**; see also Matchin & Wood, 2020 for a similar predictions figure), the effect of phrase-structure sensitivity in the IFG language fROI does not statistically differ between production and comprehension (p=0.66; **Table SI-4**), and the magnitude of the SProd>WProd effect (as well as the SComp>WComp effect) is strikingly similar between the IFG and PostTemp language fROIs (**Figure 4b**). Instead, in both the IFG language fROI and the PostTemp language fROI, the language production conditions elicit a stronger response than the language comprehension conditions (*p*s<0.02). This effect also holds for the language network as a whole (p<0.001) and is present in all language fROIs, except for the AngG fROI (**Table SI-4**).

**Figure 4.**
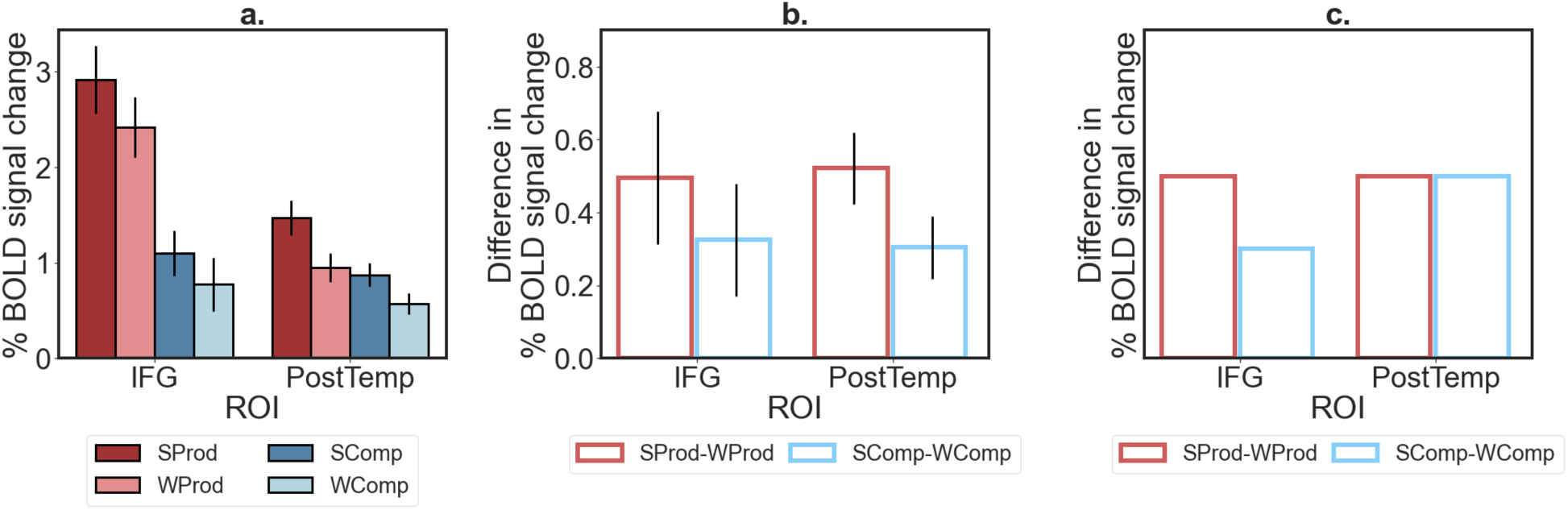
Responses of the IFG and PostTemp language fROIs during sentence and word-list production and comprehension. a. Responses of the IFG and PostTemp language fROIs to the language production and comprehension conditions: sentence production (SProd), word-list production (WProd), sentence comprehension (SComp), and word list comprehension (WComp). Error bars represent standard errors of the mean over participants. b. Observed difference in response magnitudes for the SProd–WProd effect (red bars) and the SComp–WComp effect (blue bars). c. A pattern of differences in response magnitudes for the SProd–WProd and SComp–WComp effects in the two ROIs, as predicted by Matchin & Hickok’s (2019) proposal.

Our whole-brain search for brain regions that support phrase-structure building during production (as evidenced by the SProd>WProd effect) and also respond more strongly during sentence production than sentence comprehension (as evidenced by the SProd>SComp effect) revealed 9 regions (**Figure SI-3**). However, based on the full response profiles of these fROIs, none of them appear to be selective for phrase-structure building during production. In particular, two kinds of profiles characterize these regions. Four fROIs (marked in teal in **Figure SI-3**) in the posterior inferior temporal, occipital, and cerebellar cortex respond strongly to the visual event semantic condition, suggesting that their strong and selective response to the sentence production condition (relative to the word-list production and sentence comprehension conditions) is driven by the visual and/or semantic demands associated with the processing of event pictures (see also Ivanova et al., in prep). And the remaining five fROIs (marked in yellow in **Figure SI-3**) in the left lateral and medial frontal cortex appear to overlap with the extended language-selective network, as they all show a robust response to the language localizer contrast (and no response to the MD localizer). The extended language network includes a number of cortical, as well as subcortical and cerebellar, areas outside of the left frontal and temporal cortex (e.g., Fedorenko et al., 2011; Lipkin et al., in press), including the medial frontal areas that emerge in this analysis. It is not clear why these fROIs show relatively weak responses to the sentence comprehension (SComp) condition, not strongly differing from the response to the word-list comprehension (WComp) condition (cf. the response to the language localizer contrast). One possibility is that the sentences in the SComp condition are relatively short and simple, and semantically and structurally similar to one another (Appendix A; cf. the sentences in the language localizer, which are longer and more semantically and syntactically diverse; https://evlab.mit.edu/funcloc/). In any case, the strong response to the comprehension-based localizer contrast shows that these regions are not strongly selective for phrase-structure building in language production and also support high-level linguistic comprehension.

## Discussion

We examined neural responses to cognitive demands associated with high-level language production—lexical access and phrase-structure building—in the fronto-temporal language-selective network (Fedorenko et al., 2011) using a robust precision fMRI approach. Across three experiments that employed a picture naming/description paradigm to elicit word- and sentence-level productions, we found that i) sentence production, spoken or typed, elicits a strong response across the entire language-selective network as defined by a comprehension-based localizer (Fedorenko et al., 2010); ii) the language network is sensitive to both phrase-structure building and lexical access, but phrase-structure building demands elicit a stronger and more spatially extensive effect, reliably manifesting in every language fROI; and iii) no region within the language network, or in the rest of the brain, appears to selectively support phrase-structure building in production relative to comprehension. Below, we contextualize these results in the current theoretical and empirical landscape of the field and discuss their implications.

### Ubiquitous sensitivity to sentence production across the language network

Sentence production appears to recruit the entire fronto-temporal language-selective network. In each of the six language fROIs, we observed a large and reliable response during the sentence production condition (spoken or typed) in all three fMRI experiments. Because we focused on the fROIs that have been unambiguously and selectively linked to high-level language processing, including lexico-semantic and combinatorial processes (e.g., Fedorenko et al., 2010, 2020; Bautista & Wilson, 2016), we can conclude that high-level comprehension and production draw on the same network, ruling out the possibility that the overlap is due to shared perceptual and/or motor demands. This finding aligns with earlier findings from neuroimaging studies that have relied on the group-averaging approach and reported overlap in parts of the language network (e.g., Awad et al., 2007; Menenti et al., 2011; Segaert et al., 2012; Silbert et al., 2014; Giglio et al., 2012), but goes beyond them in two ways. First, the individual-subject analytic approach ensures that the overlap between comprehension and production is not due to the artifacts of blurring nearby distinct regions, which is inherent in traditional group analyses (e.g., Nieto-Castañón & Fedorenko, 2012).

And second, this is the first study that demonstrates that responses to sentence production are robust to the output modality (speaking vs. typing). In particular, a key signature of the language network is input-modality independence during comprehension, as evidenced by similar responses across listening and reading (e.g., Fedorenko et al., 2010; Vagharchakian et al., 2012; Regev et al., 2013; Scott et al., 2017; Chen et al., 2021), as well as during visual processing in sign language comprehension (e.g., MacSweeney et al., 2008). Here, we show that the language network also exhibits output-modality-independent responses during production, across speaking and typing. This generalization across modalities demonstrates that the observed effects concern higher-level aspects of production (access of linguistic representations and utterance planning) rather than lower-level implementation parts of the production pipeline. As discussed in the Introduction, the network of areas that support articulation (e.g., Bohland & Guenther, 2006; Fedorenko & Thompson-Schill, 2014; Basilakos et al., 2018; see **Figure SI-2a** for a profile of regions in this network) is distinct from the higher-level language network examined here. The hand motor control areas associated with writing or typing linguistic utterances have been less extensively investigated, but are also distinct from the language network (e.g., Roux et al., 2009; Longcamp et al., 2014; Willet et al., 2021; **Figure SI-2b**).

The fact that every region of the language network responds during both interpretation and generation of linguistic utterances suggests that this network plausibly stores our language knowledge—mappings between forms and meanings—that are, of course, necessary for both comprehension (by evaluating the input relative to these stored representations) and production (by searching these representations for the right words/constructions). These results align well with past theoretical proposals (e.g., Strikers & Costa, 2016; Gambi & Pickering, 2017). The fact that responses to sentence production (as well as to sentence comprehension) are distributed across the language network aligns with growing evidence that this network constitutes a ‘natural kind’ in the mind and brain, working as an integrated system to solve comprehension and production (e.g., Mesulam, 1990; Blank et al. 2014; Fedorenko & Thompson-Schill, 2014; Braga et al., 2020).

### The language network supports both lexical access and phrase-structure building

As discussed in the Introduction, high-level production includes selecting the right words and constructions (lexical access) and putting them together into well-formed strings, including ordering the words and selecting the right form of each word according to the intended meaning and the structure being built (e.g., selecting the plural form of a noun, or selecting the right tense for a verb) (phrase-structure building). We found that the language network is sensitive to cognitive demands associated with both of these components of high-level production, although the response to phrase-structure building demands is generally stronger.

We used a paradigm adapted from comprehension (e.g., Friederici et al., 2000; Humphries et al., 2007; Fedorenko et al., 2010) in an effort to separate lexical access and phrase-structure building demands. In comprehension, the question of whether different brain regions in the language network support the understanding of individual word meanings vs. combinatorial (syntactic/semantic) processing has long been controversial. Based on the critical review of the literature and several additional studies, Fedorenko et al. (2020) argued that no brain region within the language network is selective for combinatorial processing over the processing of single words (see also Chee et al., 1999; Keller et al., 2001; Röder et al., 2002; Bautista & Wilson, 2016; Blank et al., 2016; see Toneva & Wehbe, 2019; Schrimpf et al., 2021; Caucheteux et al., 2021 for converging evidence from relating human neural representations to those from artificial neural network models; and see Dick et al., 2001 for earlier arguments and evidence against syntax selectivity).

Here, we asked this question for language production. The response to phrase-structure building demands (evidenced by a stronger response during sentence production than word-list production) was reliable in every language fROI. This distributed nature of phrase-structure building a) parallels the distributed effects of syntactic demands during language comprehension (e.g., Blank et al., 2016; Shain, Blank et al., 2020; Shain et al., in press), and b) aligns with evidence from aphasia, where damage to both frontal and temporal language areas and the white matter tracts connecting them can result in syntactic deficits (e.g., Kempler et al., 1991; Caplan et al., 1996; Dick et al., 2001; Wilson & Saygin, 2004; Mesulam et al., 2015; Wilson et al., 2022; see deBleser, 1987 for a discussion of earlier evidence), thus adding to the growing evidence against focal implementation of combinatorial linguistic processing. Importantly, we showed that the sentence > word-list effect in production cannot be explained by general cognitive demands. In particular, the domain-general Multiple Demand network (Duncan, 2010, 2013)—which is sensitive to effort across domains (e.g., Duncan & Owen, 2001; Fedorenko et al., 2013; Hugdahl et al., 2015; Shashidara et al., 2019; Assem et al., 2020)—responded more strongly during word-list production than sentence production, in line with a similar effect that had been reported for comprehension (e.g., Diachek, Blank, Siegelman et al., 2020).

The response to lexical access demands was positive in all six language fROIs, but generally lower than the response to phrase-structure building and only statistically reliable in three of the six fROIs. This picture differs somewhat from what has been reported for language comprehension, where lexical access demands manifest reliably in every region of the language network (e.g., Fedorenko et al., 2010; Shain et al., 2021). Why the responses to lexical access demands in production in the current study were not as robust as those previously reported for comprehension is not clear. One possibility is that the objects that we used all had relatively high-frequency names, and some of these words may have already been activated in other conditions (e.g., sentence production or sentence/word-list comprehension) due to the semantic and lexical overlap in the materials. We leave to future work to investigate responses to lexical access demands in greater detail and across a wider range of materials and paradigms. We also acknowledge that some areas outside the boundaries of the language network, as defined here, may selectively contribute to lexical access in production, as has been suggested in some patient studies (e.g., Bi et al., 2011; Mesulam et al., 2013).

Evidence from aphasia deserves a brief mention given that some patients with aphasia have been argued to exhibit a selective grammatical deficit (‘agrammatism’; see deBleser, 1987 for a historical overview). We would argue that no single study has compellingly established the existence of syntax-selective machinery (see also e.g., Badecker & Caramazza, 1985; Berndt et al., 1996; for recent claims about the existence of such machinery, see e.g., Matchin et al., 2020). Such a demonstration would require establishing that the syntactic deficit a) is present in both production and comprehension (i.e., affects the underlying syntactic representations), b) generalizes across spoken and written modalities, diverse linguistic materials, and experimental paradigms, c) cannot be explained by low-level perceptual or motor difficulties, or non-linguistic factors (e.g., executive limitations), and d) is selective relative to other aspects of language (like lexical access). In fact, some evidence exists against a syntax-selective deficit in aphasia. First, anomia (lexical retrieval difficulties) is *ubiquitous* in aphasia, including for patients with agrammatic production and/or comprehension (e.g., Goodglass & Geschwind, 1976; Blumstein, 1988; see Lu et al., 2021 for related evidence from cortical stimulation), suggesting that brain areas whose damage leads to syntactic deficits also contribute to lexical access (see also Bates & Goodman, 1997). And second, syntactic comprehension deficits and the kinds of errors that are observed in patients with expressive agrammatism have been shown to be inducible in neurotypical adults (e.g., Butterworth & Howard, 1987; Miyake et al., 1994; Blackwell & Bates, 1995), arguing against a representation-level explanation of such behaviors.

### No evidence of brain regions that are selective for phrase-structure building during language production

We do not find evidence for the existence of brain mechanisms that selectively support phrase-structure building during production relative to comprehension. Such mechanisms have been hypothesized to exist given that morpho-syntactic processes play a critical role in sentence encoding during production (cf. Swets et al., 2013; Goldberg & Ferreira, 2022) but are not strictly necessary during sentence comprehension, which is possible even when such cues are degraded or absent (e.g., Ferreira et al., 2002; Ferreira, 2003; Levy, 2008; Levy et al., 2009; Gibson et al., 2013; Ferreira & Lowder, 2016; Mahowald, Diachek et al., 2022). We evaluated a particular proposal due to Matchin & Hickok (2019), whereby the inferior frontal component of the language network is hypothesized to selectively support morpho-syntactic planning for language production (relative to comprehension). We found that effects associated with phrase-structure building demands were similar for production and comprehension in the inferior frontal (IFG) language fROI, and this pattern was similar to that in the posterior temporal (PostTemp) and other language fROIs. Instead, language regions responded overall more strongly to the production than the comprehension conditions, perhaps unsurprisingly given that production trails comprehension in development and is more challenging for adult language learners (e.g., Jakobson 1941/1968). We also searched across the brain for regions with the hypothesized profile (a selective response to phrase-structure building during production) and did not find such regions.

Taken at face value, the lack of the asymmetry in our data between the inferior frontal and posterior temporal language regions is at odds with two recent published fMRI findings. In particular, Matchin & Wood (2020) and Giglio et al. (2022) both claim to find support for Matchin & Hickok’s (2019) hypothesis. However, both studies have limitations that makes their results difficult to interpret. Matchin & Wood (2020) report a study where participants i) read, ii) read and subvocally repeated, or iii) listened to Jabberwocky phrases (e.g., “these clopes this pand”), real words (e.g., “hermit dogma”), or nonwords (e.g., “ninyo pobset”). One problematic aspect of the design is that the production condition (reading and subvocally repeating strings) does not require high-level planning: participants simply repeat the already constructed stimuli (presented to them on the screen). More importantly, however, the authors do not report the magnitudes of response to their conditions, as would be needed for evaluating the hypothesis in question. In particular, the main results figure in their paper (Figure 4) shows the *t-*statistics for the relevant contrasts, not beta weights or percent BOLD signal change values. Because *t*-values are scale-free (the ratio of the effect size to the standard error), they do not tell us about the size of the change in the BOLD signal. Instead, they provide a measure of confidence in the deflection from zero, with bigger *t-*values reflecting greater confidence in a positive effect, not a larger effect. As a result, as reported, the findings do not speak to Matchin & Hickok’s (2019) hypothesis.

Giglio et al. (2022) report a study that adapted the design from Pallier et al. (2011), who had compared neural responses to linguistic strings composed of the same number of words but varying in the syntactic structure so as to allow for shorter vs. longer composite phrases. Pallier et al. (2011) found that the neural response in the language regions increases as a function of the length of composite phrases (e.g., a string like “a girl with pigtails” elicits a stronger response than a string like “a girl a bag”). Giglio et al. developed a paradigm where on each trial, participants saw pictures of two possible agents (e.g., a girl and a woman) and were provided with the verbs (e.g., “to clap”, “to sleep”). They were then asked to either view the pictures and listen to the accompanying linguistic strings (comprehension conditions) or to generate linguistic strings (production conditions). Critically, the way the pictures were structured on the screen indicated the target syntactic structure. For example (as shown in Figure 1 in Giglio et al.’s paper), when each agent had a box around it and a verb in a box above each, participants heard— or were asked to generate—strings like “a girl claps, a woman sleeps” (two clauses, each two content words long), but when the agents and the verbs were all within the same box, participants heard, or were asked to generate, strings like “a girl hears that a woman claps” (a clause that is four content words long). The authors conceptually replicated Pallier et al.’s (2011) results of stronger neural responses to longer phrases in their comprehension conditions (see also Shain et al., 2021). They also found a similar pattern in their production conditions. Of most relevance to the current investigation, they report similar slopes for these structural effects for comprehension and production in their middle temporal gyrus (MTG) ROI, and a higher slope for production in their inferior frontal gyrus (IFG) ROI. They take this pattern as evidence for a greater role of the IFG ROI in structure building during language production, compared to comprehension. In the Giglio et al.’s study, the interpretation difficulty results from how the ROIs are defined. In particular, the authors use large anatomical masks (Figure 4d in their paper): one encompasses the entire middle temporal gyrus, and the other encompasses the inferior frontal gyrus. This is problematic because in the inferior frontal cortex, language-selective areas lay adjacent to the areas of the Multiple Demand (MD) network, which is functionally distinct from the language network (Fedorenko et al., 2012b see Fedorenko & Blank, 2020 for review). Because MD areas are robustly sensitive to task difficulty across domains (e.g., Duncan & Owen, 2000; Fedorenko et al., 2013; Shashidara et al., 2019; Assem et al., 2020) and because the structurally more complex production conditions are more difficult (as evidenced by the behavioral data in Figures 2 and 3a in their paper), the inclusion of parts of the MD cortex in the region of interest effectively guarantees the pattern observed for the IFG ROI. (The MTG ROI also includes some non-language selective cortex given what we know about the topography of individual language areas (e.g., Fedorenko et al., 2010; Lipkin et al., in press), but it does not include areas that are sensitive to general cognitive effort.)

We acknowledge the possibility that the lack of selectivity for morpho-syntactic encoding in production in our data may have to do with the fact that many participants relied on ‘headlinese’ (SI-1) at least some of the time (i.e., saying or typing “girl smelling a flower” instead of “A girl is smelling a flower”). Headlinese has its own set of syntactic constraints (e.g., Halliday, 1967; Mårdh, 1980; van Dijk 1988), and whether morpho-syntactic demands that are associated with the production of headlinese-style utterances are lower compared to the production of complete well-formed sentences is an empirical question that, to the best of our knowledge, has not been tackled before. But even if producing headlinese is easier with respect to morpho-syntactic encoding, it seems valuable to investigate sentence production under as natural conditions as possible. During everyday communication, we rarely speak in complete sentences, and if many/most participants produce headlinese-style responses when asked to describe pictures of events, then it seems worthwhile to understand the cognitive and neural infrastructure supporting this behavior.

The data pattern we observe here, along with the findings reported in past studies of language comprehension (e.g., Fedorenko et al., 2010, 2012a, 2020; Pallier et al., 2011; Blank et al., 2016; Shain, Blank et al., 2020)—whereby phrase-structure building demands elicit strong responses across the language network during both comprehension and production—aligns with recent evidence from Shain et al. (in press). Shain and colleagues report a large-scale fMRI study that suggests that language comprehension involves computationally demanding word-by-word structure building operations even when participants passively listen to naturalistic stories. Thus, although comprehension is possible in many cases where morpho-syntactic cues are degraded or absent (e.g., Ferreira et al., 2002; Ferreira, 2003; Levy, 2008; Levy et al., 2009; Gibson et al., 2013; Ferreira & Lowder, 2016; Mollica et al., 2020; Mahowald, Diachek et al., 2022), it appears that rich syntactic structures are nevertheless always computed, contrary to arguments that human language processing is mostly approximate and shallow (e.g., Frank & Bod, 2011).

In conclusion, we have shown that the language-selective network, which supports comprehension across modalities, also supports sentence production during speaking and typing. Similar to the strong integration between the processing of word meanings and combinatorial processing that has been observed for language comprehension (e.g., Fedorenko et al., 2020), we found that the language network is sensitive to both the demands associated with phrase-structure building and those associated with lexical access during language production, although the effects of phrase-structure building are more pronounced and reliably present in every language region. Finally, contra prior hypotheses, we did not find evidence of brain areas that are selective for phrase-structure building during production relative to comprehension; instead, sentence production appears to pose a higher cost to the language network than sentence comprehension. These results support the idea that the language network stores integrated linguistic knowledge, from phonotactic regularities, to morphological schemas, to words, to constructions, and this knowledge is accessed during both decoding and encoding of linguistic messages.

## Acknowledgements

We would like to acknowledge the Athinoula A. Martinos Imaging Center at the McGovern Institute for Brain Research at MIT, and its support team. We thank former and current EvLab members (especially Josef Affourtit and Alvincé Pongos) for their help with data collection, Rachel Ryskin for extensive discussions of the relationship between comprehension and production, and the audience at the 2020 Neurobiology of Language conference and members of the Fedorenko and Gibson labs for helpful comments and discussions.

