## Supplementary Information for "Precision fMRI reveals that the language-selective network supports both phrase-structure building and lexical access during language production"

* Equal contributors

### Supplementary Tables

| Language Localizer | The experiments that used that language localizer version and number of subjects in each | Trial Structure | # trials & blocks | Task | Total duration |
|---|---|---|---|---|---|
| **Version 1** | Expt 1 = 29,<br>Expt 2 = 6 | 100ms trial-initial fixation; 12 words/nonwords presented for 450ms each; 400ms button press; 100ms trial-final fixation | 3 trials per block, 8 blocks per condition,<br>16 blocks total | Button press | 5 minutes, 58 seconds |
| **Version 2** | Expt 2 = 4 | 300ms trial-initial fixation; 12 words/nonwords presented for 350 ms each; 1000 ms probe; 500 ms trial-final fixation | 3 trials per block, **8 blocks** per condition,<br>16 blocks total | Memory probe | 6 minutes, 18 seconds |
| **Version 3** | Expt 2 = 2 | 300ms trial-initial fixation; 12 words/nonwords presented for 350 ms each; 1000 ms probe; 500 ms trial-final fixation | 3 trials per block, **6 blocks** per condition,<br>18 blocks total | Memory probe | 6 minutes, 36 seconds |

**Table SI-1. Language localizer details for all experiments.** (As described in the main text, participants in Experiment 3 were a subset of the participants in Experiment 1.)

| fROI | Experiment 2 | | Experiment 3 | | | |
|---|---|---|---|---|---|---|
| | SProd vs. fixation | SProd vs. Nonwords | SProd (typed) vs. fixation | SProd (typed) vs. Nonwords | SProd (typed) vs. NProd (typed) | SProd (typed) vs. VisEvSem |
| **Language network** | **d = 1.731**<br>**p < 0.001** | **d = 1.587**<br>**p < 0.001** | **d = 1.027**<br>**p < 0.001** | **d = 0.498**<br>**p = 0.003** | **d = 0.705**<br>**p < 0.001** | **d = 0.576**<br>**p < 0.001** |
| IFGorb | $d = 1.656$<br>$p = 0.012$ | $d = 2.300$<br>$p = 0.003$ | $d = 1.507$<br>$p = 0.018$ | $d = 1.263$<br>$p = 0.121$ | $d = 3.615$<br>$p < 0.001$ | $d = 1.603$<br>$p = 0.025$ |
| IFG | $d = 2.449$<br>$p = 0.001$ | $d = 2.452$<br>$p = 0.003$ | $d = 1.771$<br>$p = 0.018$ | $d = 1.283$<br>$p = 0.121$ | $d = 1.957$<br>$p = 0.006$ | $d = 1.650$<br>$p = 0.025$ |
| MFG | $d = 3.115$<br>$p < 0.001$ | $d = 3.300$<br>$p = 0.001$ | $d = 1.494$<br>$p = 0.018$ | $d = 0.361$<br>$p = 0.631$ | $d = 1.713$<br>$p = 0.010$ | $d = 1.492$<br>$p = 0.028$ |
| AntTemp | $d = 2.105$<br>$p = 0.003$ | $d = 2.519$<br>$p = 0.003$ | $d = 0.561$<br>$p = 0.198$ | $d = 0.274$<br>$p = 0.631$ | $d = 2.835$<br>$p < 0.001$ | $d = 0.326$<br>$p = 0.680$ |
| PostTemp | $d = 2.434$<br>$p = 0.001$ | $d = 2.410$<br>$p = 0.003$ | $d = 1.443$<br>$p = 0.018$ | $d = 0.488$<br>$p = 0.631$ | $d = 3.104$<br>$p < 0.001$ | $d = 1.764$<br>$p = 0.025$ |
| AngG | $d = 1.393$<br>$p = 0.025$ | $d = 1.575$<br>$p = 0.003$ | $d = 0.015$<br>$p = 0.969$ | $d = 0.006$<br>$p = 0.992$ | $d = 1.625$<br>$p = 0.012$ | $d = -0.077$<br>$p = 0.892$ |

**Table SI-2. Responses in the language network to spoken and typed sentence production.** Effect sizes (Cohen's *d*) and estimated *p*-values for the effect of the spoken SProd condition (relative to different baselines) in linear mixed-effects regression models in Experiments 2 and 3 (see Analyses, Q1; see **Table 1** for the responses in Experiment 1). Models were fit to perform pairwise comparisons of sentence production (SProd) vs. each of the following conditions: fixation, nonword comprehension (Nonwords; from the language localizer), nonword production (NProd), and visual event semantics processing (VisEvSem). The results are shown averaged across the language network (top row), as well as at the level of individual functional ROIs in the language network (bottom six rows; FDR corrected). Green cells highlight significance at $p<0.05$ in the predicted direction.

| fROI | Experiment 2 | Experiment 3 | |
|---|---|---|---|
| | SProd vs. WProd | SProd (typed) vs. WProd (typed) | WProd (typed) vs. NProd (typed) |
| **Language network** | **d = 0.609**<br>**p < 0.001** | **d = 0.830**<br>**p < 0.001** | **d = −0.105**<br>**p = 0.524** |
| IFGorb | *d* = 2.727<br>*p* = 0.002 | *d* = 2.966<br>*p* < 0.001 | *d* = −0.150<br>*p* = 0.825 |
| IFG | *d* = 1.003<br>*p* = 0.124 | *d* = 1.570<br>*p* = 0.014 | *d* = −0.125<br>*p* = 0.825 |
| MFG | *d* = 1.158<br>*p* = 0.097 | *d* = 1.942<br>*p* = 0.005 | *d* = 0.148<br>*p* = 0.825 |
| AntTemp | *d* = 2.251<br>*p* = 0.005 | *d* = 2.988<br>*p* < 0.001 | *d* = −1.881<br>*p* = 0.029 |
| PostTemp | *d* = 2.816<br>*p* = 0.002 | *d* = 3.693<br>*p* < 0.001 | *d* = −0.831<br>*p* = 0.378 |
| AngG | *d* = 4.396<br>*p* < 0.001 | *d* = 2.718<br>*p* < 0.001 | *d* = −0.769<br>*p* = 0.378 |

**Table SI-3. Responses in the language network to phrase-structure building and lexical access.** Effect sizes (Cohen's *d*) and estimated *p*-values for the effects associated with phrase-structure building and lexical access in linear mixed-effects regression models in Experiments 2 and 3 (see Analyses, Q2; see **Table 2** for the responses in Experiment 1). Models were fit to perform pairwise comparisons of sentence production (SProd) vs. word-list production (WProd), and word-list production vs. nonword production (NProd), for both spoken (Experiment 2) and typed (Experiment 3) modalities. The results are shown averaged across the language network (top row), as well as at the level of individual functional ROIs in the language network (bottom six rows; FDR corrected). Green cells highlight significance at $p<0.05$ in the predicted direction.

| fROI | Experiment 1 | | |
|---|---|---|---|
| | Production vs. Comprehension | Sentences vs. Word lists | Interaction |
| **Language network** | **d = 0.564**<br>**p < 0.001** | **d = 0.183**<br>**p = 0.019** | **d = 0.136**<br>**p = 0.081** |
| IFGorb | $d$ = 0.977<br>$p$ < 0.001 | $d$ = 0.123<br>$p$ = 0.659 | $d$ = 0.376<br>$p$ = 0.159 |
| IFG | $d$ = 1.617<br>$p$ < 0.001 | $d$ = 0.320<br>$p$ = 0.240 | $d$ = 0.119<br>$p$ = 0.659 |
| MFG | $d$ = 1.459<br>$p$ < 0.001 | $d$ = 0.270<br>$p$ = 0.303 | $d$ = 0.086<br>$p$ = 0.694 |
| AntTemp | $d$ = 0.682<br>$p$ = 0.011 | $d$ = 0.478<br>$p$ = 0.062 | $d$ = 0.307<br>$p$ = 0.244 |
| PostTemp | $d$ = 0.634<br>$p$ = 0.017 | $d$ = 0.508<br>$p$ = 0.051 | $d$ = 0.257<br>$p$ = 0.312 |
| AngG | $d$ = −0.101<br>$p$ = 0.683 | $d$ = 0.559<br>$p$ = 0.036 | $d$ = 0.507<br>$p$ = 0.051 |

**Table SI-4. Responses in the language network to sentence and word-list production and comprehension.** Effect sizes (Cohen's *d*) and estimated *p*-values for the main effects of task (production vs. comprehension), stimulus (sentences vs. word lists), and the interaction between them in linear mixed-effects regression models for data in Experiment 1 (see Analyses, Q3). Models were fit to perform pairwise comparisons of language production (SProd, WProd) vs. comprehension (SComp, WComp), sentences (SProd, SComp) vs. word lists (WProd, WComp), and the interaction between them. The results are shown averaged across the language network (top row), as well as at the level of individual functional ROIs in the language network (bottom six rows; FDR corrected).

**Supplementary Figures**

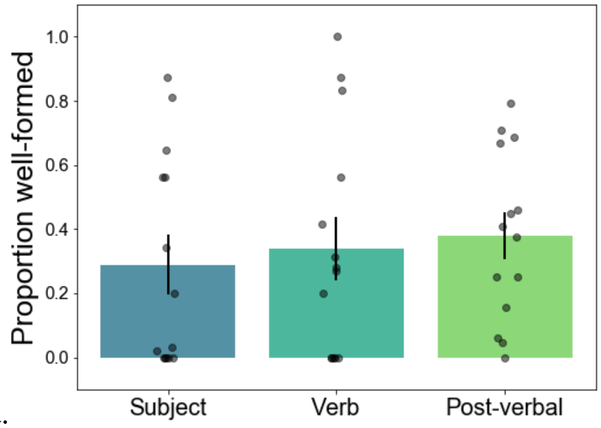

a.

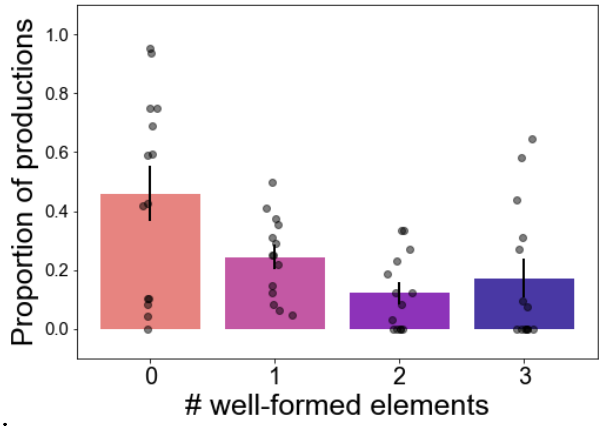

b.

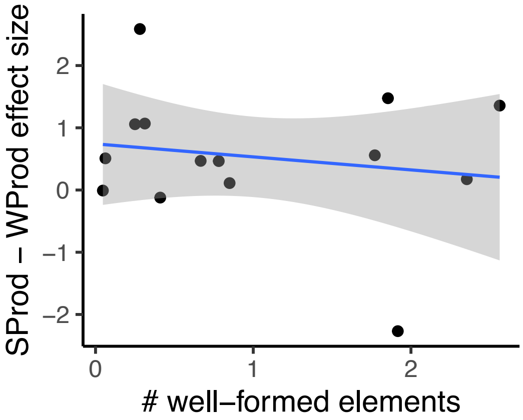

c.

**Figure SI-1. Details on the sentence productions in Experiment 3.** We examined the sentence productions (typed responses; spoken productions were not recorded). **a.** Mean proportions of grammatically well-formed (according to the rules of standard English grammar; cf. headline-style syntax) subject phrases, verb phrases, and post-verbal elements. For the subjects and post-verbal elements, we counted a phrase as well-formed if it included a determiner, and for verb phrases, we counted a phrase as well-formed if it used a tense form. Here, and in panel b, error bars represent standard errors of the mean over participants; individual dots represent participants. **b.** Mean proportion of productions with 0, 1, 2, or 3 well-formed elements. **c.** The relationship between the magnitude of the SProd>WProd effect (i.e., the contrast that targets phrase-structure building demands; the effect is averaged across the language network, and the dots correspond to participants) and the average number of well-formed elements per utterance (note that participants varied substantially, with some producing complete sentences most of the time, and others exclusively relying on ‘headlinese’). No relationship was found (Pearson r=-0.17, p=0.56).

a.

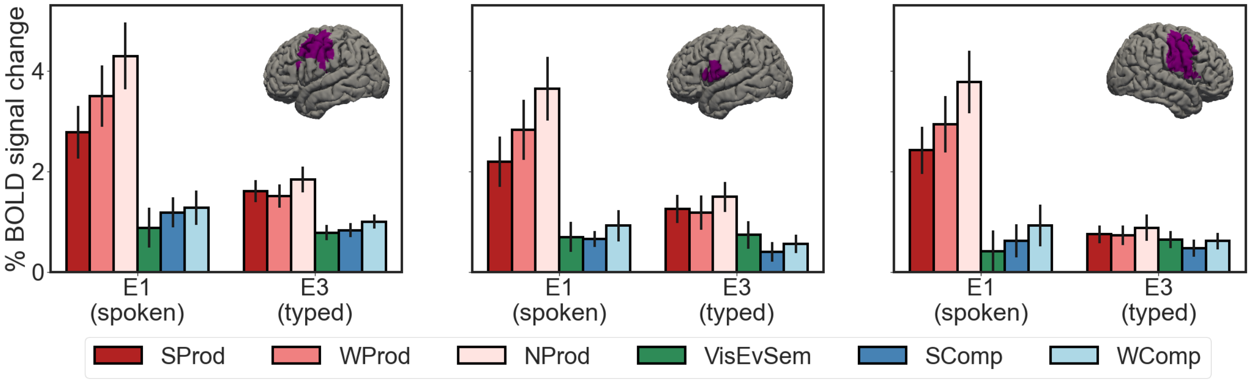


b.

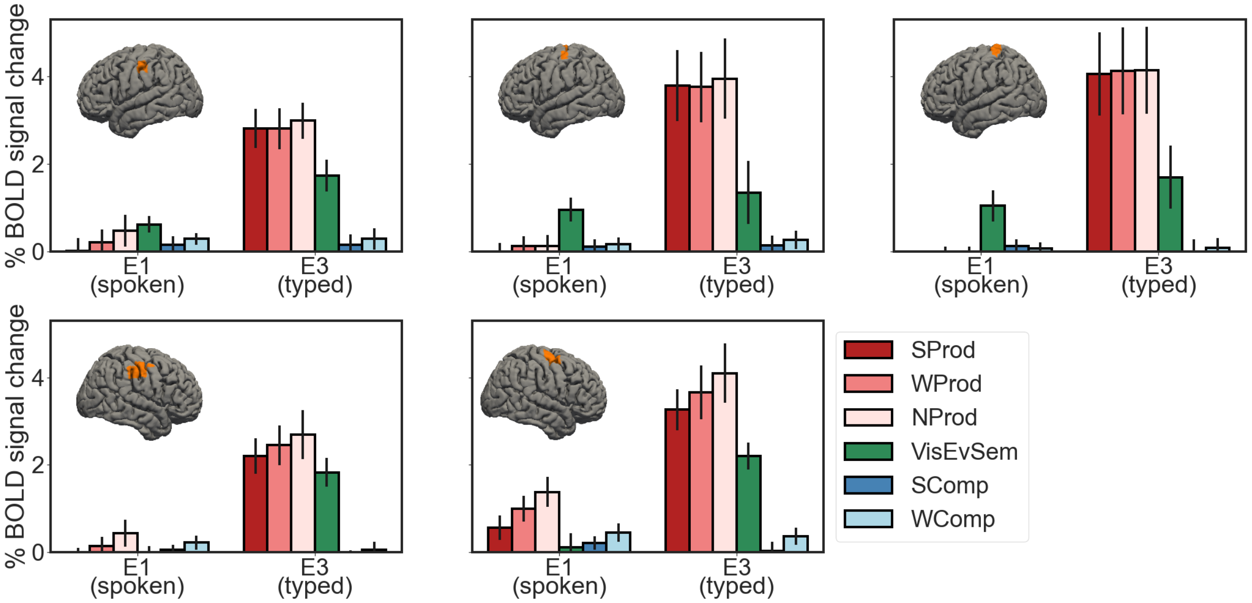


**Figure SI-2. Response profiles of select articulation-selective (a) and typing-selective (b) areas.** We performed a whole-brain group-constrained subject-specific (GSS) analysis (Fedorenko et al., 2010) for the nonword-list production (NProd) > Fixation contrast, using data from subjects who completed both the spoken and typed production (n=14). Experiment 1 (spoken production) was used to search for articulation-responsive areas and data from Experiment 3 (typed production) was used to search for typing-responsive areas. In particular, for each participant, we thresholded the whole-brain map for the relevant (spoken or typed) NProd>Fixation contrast at $p<0.001$ (uncorrected level). Each participant's map was then binarized, with 1s corresponding to voxels that show a reliable effect, and 0s otherwise. These individual maps were then overlaid in the common space, and watershed parcellation was performed, as described in Fedorenko et al. (2010), to search for areas that would contain supra-threshold voxels in at least half of the participants. The resulting regions (parcels) were then used to define the individual fROIs using the same contrast (NProd(spoken/typed)>Fix) by selecting the top 10% of voxels based on the *t*-values for the relevant contrast. In this way, a fROI was defined in every participant. Finally, we estimated the responses to the critical conditions (SProd, WProd, NProd, VisEvSem, SComp, and WComp) in these individually defined fROIs (to estimate the response to the NProd condition, across-runs cross-validation was used to ensure independence; as described in Methods for estimating the responses to the language and MD localizer conditions in the language and MD fROIs, respectively). **a.** Responses to the critical conditions in Experiments 1 and 3 for three sample articulation-responsive functional ROIs (parcels used to define these are shown as brain insets). **b.** Responses to the critical conditions in Experiments 1 and 3, for five sample typing-responsive functional ROIs. As can be seen from the response profiles, these areas show the expected selectivity for spoken (figures in a) and typed (figures in b) language production. Unlike the language fROIs examined in the main text, these regions respond similarly strongly to language production regardless of the content (sentences, word lists, or nonword lists), as expected given that lower-level articulatory/hand-motor demands are similar between these conditions. (Note that the relatively strong responses to the VisEvSem condition are likely due to the fact that participants were using their fingers to perform the button press.)

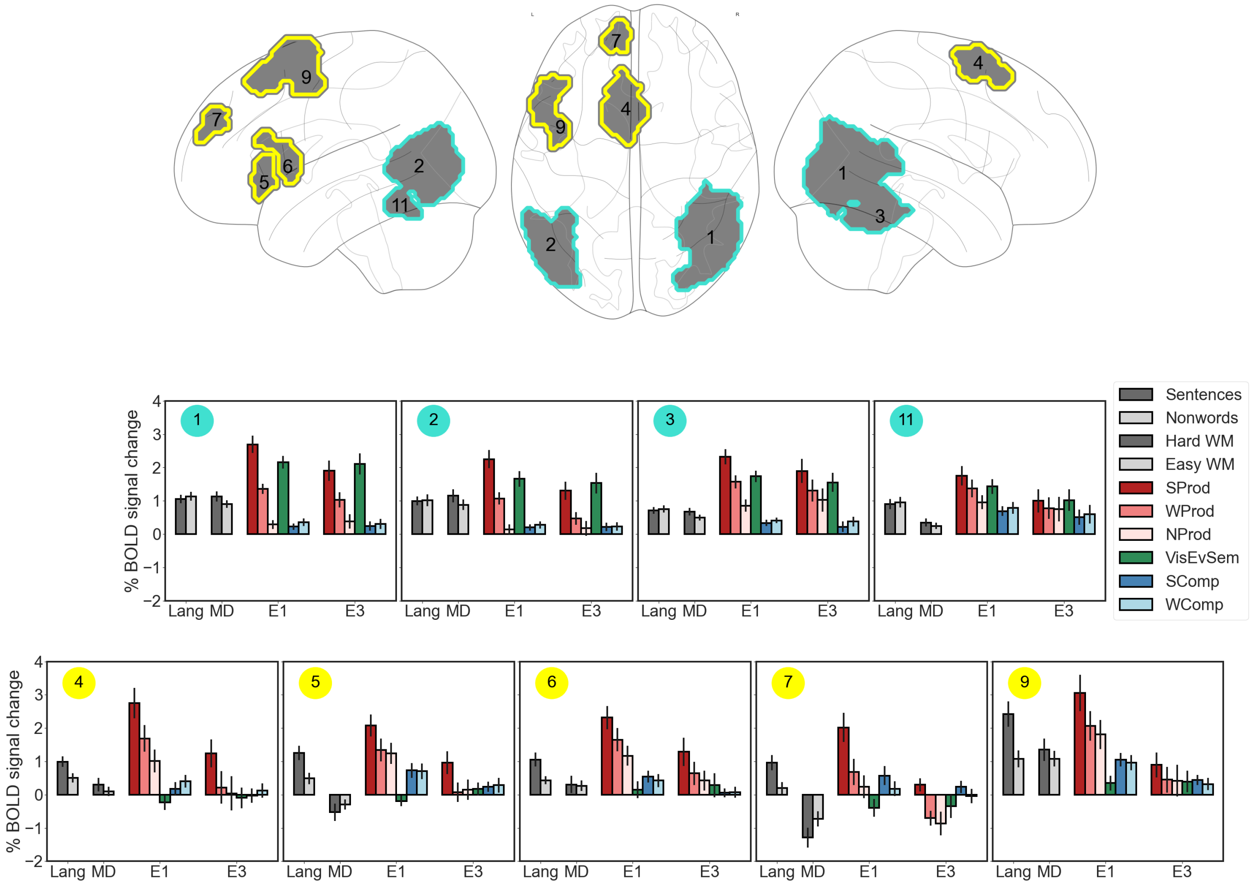


**Figure SI-3.** Top row: Brain regions that emerged when we searched across the brain for areas that selectively support phrase-structure building in production relative to comprehension. To search for such areas, we performed a whole-brain group-constrained subject-specific (GSS) analysis (Fedorenko et al., 2010) on the data from Experiment 1. We used a conjunction of the following two contrasts: i) sentence production > word-list production (SProd>WProd), and ii) sentence production > sentence comprehension (SProd>SComp) (see Q3 in Methods). Nine regions were recovered that contained supra-threshold voxels in the majority of the participants (range: 0.59-0.97) and that replicable (as assessed using across-runs cross-validation) SProd>WProd (ps< 0.0007) and SProd>SComp (ps< $10^{-8}$) effects (see below for the anatomical characterization of the parcels). The parcels are grouped (by color: teal and yellow) into two sets that correspond to the two kinds of profiles we observed (see below).

Middle and bottom rows: The functional response profiles of the fROIs that were defined in individual participants within the parcels shown in the top row (see Methods) with respect to the data from the two localizer experiments and critical Experiments 1 and 3. The fROI numbers (circled in the top left corner of each bar graph) correspond to the parcel numbers in the top row. The fROIs are grouped into two sets that correspond to the two kinds of profiles we observed: four fROIs (marked in teal, middle row) in the posterior inferior temporal and occipital cortex that respond strongly to the visual event semantic condition, suggesting that their strong and selective response to the sentence production condition (relative to word-list production and sentence comprehension conditions) is driven by the visual and/or semantic demands associated with the processing of event pictures; and the remaining five fROIs (marked in yellow, bottom row) in the left lateral and medial frontal cortex appear to overlap with the extended language-selective network, as they all show a robust response to the language localizer contrast (and no response to the MD localizer). Thus, none of these candidate areas showed selectivity for sentence production based on their full response profile.

***Anatomical characterization of the parcels:***

Below, we provide information on the overlap between each parcel and anatomical areas as defined in the Harvard-Oxford atlas (Makris et al., 2006; Frazier et al., 2005; Desikan et al., 2006; Goldstein et al., 2007). We list all anatomical areas that overlapped with each target parcel by at least 5%.

**Parcel 1 (4,228 voxels):**
1767 voxels (42%) covering 86% of iLOC r (Lateral Occipital Cortex, inferior division Right)
614 voxels (15%) covering 53% of toMTG r (Middle Temporal Gyrus, temporooccipital part Right)
526 voxels (12%) covering 11% of sLOC r (Lateral Occipital Cortex, superior division Right)
194 voxels (5%) covering 22% of OFusG r (Occipital Fusiform Gyrus Right)

**Parcel 2 (3,118 voxels):**
1487 voxels (48%) covering 73% of iLOC l (Lateral Occipital Cortex, inferior division Left)
509 voxels (16%) covering 10% of sLOC l (Lateral Occipital Cortex, superior division Left)
490 voxels (16%) covering 57% of toMTG l (Middle Temporal Gyrus, temporooccipital part Left)

**Parcel 3 (1,472 voxels):**
466 voxels (32%) covering 57% of TOFusC r (Temporal Occipital Fusiform Cortex Right)
360 voxels (24%) covering 14% of Cereb1 r (Cerebellum Crus1 Right)
265 voxels (18%) covering 17% of Cereb6 r (Cerebellum 6 Right)
178 voxels (12%) covering 23% of toITG r (Inferior Temporal Gyrus, temporooccipital part Right)

**Parcel 11 (299 voxels):**
167 voxels (56%) covering 26% of TOFusC l (Temporal Occipital Fusiform Cortex Left)
95 voxels (32%) covering 14% of toITG l (Inferior Temporal Gyrus, temporooccipital part Left)
25 voxels (8%) covering 1% of Cereb1 l (Cerebellum Crus1 Left)

**Parcel 4 (1,570 voxels):**
790 voxels (50%) covering 28% of SFG l (Superior Frontal Gyrus Left)
185 voxels (12%) covering 29% of SMA L(Juxtapositional Lobule Cortex -formerly Supplementary Motor Cortex- Left)
97 voxels (6%) covering 4% of SFG r (Superior Frontal Gyrus Right)

**Parcel 5 (392 voxels):**
200 voxels (51%) covering 31% of IFG tri l (Inferior Frontal Gyrus, pars triangularis Left)
118 voxels (30%) covering 7% of FOrb l (Frontal Orbital Cortex Left)
43 voxels (11%) covering 12% of FO l (Frontal Operculum Cortex Left)
31 voxels (8%) in other / unlabeled areas

**Parcel 6 (489 voxels):**
326 voxels (67%) covering 43% of IFG oper l (Inferior Frontal Gyrus, pars opercularis Left)
56 voxels (11%) covering 9% of IFG tri l (Inferior Frontal Gyrus, pars triangularis Left)
37 voxels (8%) covering 1% of PreCG l (Precentral Gyrus Left)

**Parcel 7 (327 voxels):**
256 voxels (78%) covering 4% of FP l (Frontal Pole Left)
34 voxels (10%) covering 1% of SFG l (Superior Frontal Gyrus Left)
37 voxels (11%) in other / unlabeled areas

**Parcel 9 (250 voxels):**
137 voxels (55%) covering 5% of MidFG l (Middle Frontal Gyrus Left)
113 voxels (45%) covering 3% of PreCG l (Precentral Gyrus Left)

**Appendix A: Stimuli**

All materials are available on GitHub (https://github.com/jennhu/LanguageProduction).
The stimuli used in the SComp, WComp, and NProd conditions are also provided below for convenience:

Sentences (SComp condition):

A girl is digging sand at the beach.
A woman brushes another woman's hair.
A young girl is petting a deer.
A little girl is playing with dolls.
A man is sleeping on a bench.
A toddler is holding a toothbrush.
A boy is brushing his teeth.
Two doctors are performing surgery.
A man is jumping over a hurdle.
A baby is crawling on the floor.
A baby is sitting in a suitcase.
A young girl is kicking a ball.
A boy is looking through a telescope.
A girl is playing with a toy train.
A little boy is opening a present.
A man is taking a shower.
A woman is reading a map.
A baby is yawning.
A man is running on a treadmill.
A little girl is blowing bubbles.
A bartender is pouring a drink.
A woman is reading a book.
A man is carrying a canoe.
A girl is coloring with crayons.
A man is sweeping the sidewalk.
A woman is tying her shoe on a soccer field.
Two girls are washing a dog.
A little girl is holding a snail.
A man is mowing the lawn.
A boy is sitting in a tree.
A woman is talking on a cell phone.
A man is throwing an axe.
A woman is cutting a man's hair.
A man is sweeping the steps.
A little girl is walking on a treadmill.
A woman is putting on lipstick.
Two men are shaving their heads.
A girl is swinging on a swing.

A group of people are doing yoga on a beach.
A woman is blowdrying her hair.
A man is bowling.
A woman is eating noodles.
A young girl is climbing a fence.
A woman is blowing on a dandelion.
A boy is using a magnifying glass to look at a rock.
A group of dogs are playing in a pool.
A man is playing the flute.
A man is painting a column.
A man is eating an ice cream cone.
A young boy is playing in the sand.
A person is skiing down a steep slope.
A man is surfing.
A man is jumping off a cliff into the water.
A boy is holding up a fish he caught.
Three girls are playing jump rope.
Two kids are shoveling snow.
A woman is riding a horse.
A man is looking through a microscope.
A man is sitting on his skateboard.
A woman is sewing on her sewing machine.
Chefs are preparing food in a kitchen.
A man is rock climbing.
A woman is putting on a boot.
A young boy is sleeping.
A boy is watering a plant.
A group of people are playing poker.
A man is digging a hole.
A man is reading the newspaper.
Two older men are walking on the beach.
A woman is carrying a bowl of fruit on her head.
A young girl is covering her eyes.
A woman is holding a chicken.
A man is smoking a pipe.
A man is reading a book.
A girl is hula-hooping.
A young girl is sitting on a railing.
A little boy is holding two large cucumbers.
A man is playing the accordion.
A woman is holding a puppy.
A man is meditating in a park.
A woman is sitting on the subway.
A man is pushing a cart full of grocery bags.
A man is playing pool.
A woman is holding a stop sign.

A man is playing the saxophone.
A man is pushing a couch down a sidewalk.
A man is ironing a shirt.
A man is juggling three balls.
A man is blowing bubbles.
A woman is smiling.
A man is juggling balls.
A man is vacuuming.
A woman is taking a picture of a plant.
A man is chopping wood.
A woman is painting her nails.
A man is washing windows.
A woman is playing the harp.
A woman is diving to catch a frisbee.
A little boy is pointing at something.
A group of people are hiking up a mountain.
A man is shaving his face.
A choir is singing.
A man is holding a newborn baby.
A little girl is eating a strawberry.
A woman is taking a picture.
Two boys are jumping on a trampoline.
A woman is blowing leaves with a leaf blower.
A young child is eating an apple.
A dog is swimming in a pool.
A baby is crying.
A woman is roller skating.
A girl is playing the violin.
A man is playing guitar.
A woman is grooming a horse.
A woman is jogging.
A boy is playing the piano.
Two girls are arm-wrestling.
Two men are playing checkers.
A man is sitting on a bench reading the newspaper.
A boy is holding a bird in his hands.
A woman is washing the dishes.
A little girl is jumping into a pool.
A man is painting a castle.
A little girl is smelling a flower.
A woman is fishing.
A man is painting a mural.
A young girl is rock climbing.
A man is writing on a chalkboard.

Word lists (WComp condition):

paintbrush, shower, pool table
blowdryer, cucumbers, suitcase, broom
cucumber, guitar, brush
saxophone, couch, jumprope, lawn
hand, fruit, juggling balls, railing
puppy, chalkboard, axe
piano, frisbee, stairs, apple
shirt, puppy, mural, leaves
hole, deer, swimming pool
cocktail, doll, rollerskates
eye, shovel
fence, sewing machine, suitcase
castle, paintbrush, strawberry
dishes, plant, violin, leaves
hand, rock, treadmill, broom
toothbrush, water bottle, camera
sidewalk, shirt, ladder
canoe, toy train, dog
lawnmower, sponge
bowl, pipe, flute
shoe, bench, horse
noodle, mountain, plant
bird, apple, skateboard
stairs, ice cream
bird, razor
harp, microscope
lipstick, dog, mural, hair
skis, grocery bag, flower
lawnmower, cucumbers
crayon, hurdle, grocery bag
wave, train, railing
horse, telescope, shopping cart
wood, tooth, dandelion
dog, column, wood, hair
broom, hurdle, beach, cellphone
boot, pillow, telescope
ball, trampoline, pillow, violin
chalk, bench, microscope, chicken
violin, iron, newspaper, hand
chalk, stairs, fish
shopping cart, piano, crayon
fence, skateboard, horse, trampoline
cliff, bird, sponge
sand, bubble, accordion

skateboard, rock, strawberry
couch, bowl, swimming pool, swing set
swimming pool, baby, sidewalk
razor, plant, hair
pipe, chalk, dishrack, hairbrush
shower, mountain, pillow
column, dolls
bubble, arm
swimming pool, toothbrush, snail, fence
suitcase, iron, trampoline
cellphone, deer, cucumber
rollerskates, shovel, cliff, water bottle
arm, deer, bubble
noodle, toothbrush, book, lawn
saxophone, toy train, hairbrush
chair, cocktail, deer
treadmill, chalk
hole, scissors, noodle
fruit, bowling ball
bowling ball, tree, lipstick
water bottle, iron, card, hand
cards, vacuum, shopping cart
cards, camera
snail, ice cream, guitar
flute, paintbrush, chicken
magnifying glass, bird, ski
leaves, pipe, column
beach, arm, rock, razor
toy train, accordion, chicken
magnifying glass, lawn
hole, magnifying glass
tree, checkers, treadmill
rollerskates, eye, mountain
beach, pool table, grocery bag, map
tooth, shoe
jumprope, book, baby, flower
snow, cliff, strawberry
toothbrush, cellphone, map
bubble, hand, canoe
accordion, ladder, apple
boot, checkers, telescope
boot, tree
hurdle, brush, newspaper
dandelion, book, snail, gift
sidewalk, frisbee, vacuum, harp
dishrack, hula hoop, snow, newspaper

juggling balls, wave
saxophone, camera, bowl
lipstick, flute, castle, ball
book, wood, ball, saxophone
bowling ball, house plant, pillow, eye
snow, tooth
frisbee, shopping cart, eye
ice cream, jumprope, house plant
flower, train
shoe, telescope, noodle, skateboard
castle, dandelion, violin
chalkboard, dolls
hula hoop, shirt
window, shower, chalkboard
blowdryer, dishes
window, hairbrush
guitar, house, dishes
flute, sidewalk, cocktail, railing
couch, fish
gift, hole
fish, swing set
blowdryer, ladder, baby
axe, hand, shopping cart
lawnmower, vacuum, jumprope
newspaper, window
microscope, brush
scissors, castle, bench
chair, skis
swing set, hula hoop, shovel, hand
pipe, canoe
train, juggling balls
gift, dishrack, crayon
fruit, puppy
hula hoop, house plant, sponge, map
sand, sewing machine, chair
axe, cocktail, sewing machine
harp, strawberry, sand
scissors, beach, pool table, piano

<u>Nonword lists (NProd condition):</u>

kilch, phroose, whalse, thrarce
drautch, blourse, shramn, rhonk
shruke, stresk, snegg, wrult
spreeth, shradge, phraw, plonch
whoob, glure, shrutch, shreal

fruzz, phreep, shry, splass
twack, gwurp, twont, sprath
smon, sproe, phlish, scroat
trolf, fluint, plybe, spruite
phlooge, smu, gwoam, trooch
gletch, snolf, swirr, snebe
prauce, skourge, zorch, dwoal
shraim, thuice, skise, snarsh
throoge, twirth, plarsh, gweck
phroal, glett, blez, sweach
thwuip, whulp, frolve, yerb
sproth, scrif, flurp, blait
phloul, thrab, twarse, twaun
praught, gwawk, sneash, pruice
spreuth, phlaise, swirge, shrod
plig, swef, bleant, dwoss
phran, slasque, thwess, skole
splauge, scrank, pleeth, drys
slurch, snerth, thwange, drete
thwoque, smesh, phlieve, thwemp
threll, frung, vesk, thwee
gnant, thoule, phruiche, sploat
thoff, snith, froop, phimp
smise, splegg, scroun, gnolt
swulge, flomp, splynch, sprep
phrybe, sprelm, splure, prulge
scruite, knulf, yoove, thwoathe
frisp, shrick, phluice, thwin
blolt, sloal, sneaf, tworce
swoile, presh, dwesh, skabe
gnirp, sploc, scrinch, swelm
sprash, sproat, streve, blaub
bluin, stralf, twum, thrulch
glerk, juth, rhalp, shriece
jalp, thwoove, sprusp, scrile
zooth, glanc, phlourge, phlidge
glisque, troar, screague, glowse
plisc, snult, thutch, gwerf
splunch, spleat, therge, drelt
drooch, wrilk, splisk, trome
dwurb, smult, zoute, drealt
scrusk, sprike, flarge, twuiff
thrilt, proil, vodge, splooge
smeague, flepe, blien, thrutt
proat, smawn, squark, twag
phlin, stridge, skus, dwib

plogue, scroot, knarp, sweuth
plouth, strauve, yenge, thwurch
vilge, snalp, snyle, shraff
blorque, ghimp, phrauce, trume
gweague, drus, flimn, shruv
splybe, rhunge, viff, splieve
sloin, dwalf, thralc, fralk
strutch, sprete, spreace, sleff
knoosh, spliege, thaif, thoz
slez, twaul, gwug, phrief
dwike, skeave, splome, tralse
blorce, stroff, threpe, thorge
swulb, scroise, smirth, thisp
